# Autism genes converge on microtubule biology and RNA-binding proteins during excitatory neurogenesis

**DOI:** 10.1101/2023.12.22.573108

**Authors:** Nawei Sun, Noam Teyssier, Belinda Wang, Sam Drake, Meghan Seyler, Yefim Zaltsman, Amanda Everitt, Nia Teerikorpi, Helen Rankin Willsey, Hani Goodarzi, Ruilin Tian, Martin Kampmann, A. Jeremy Willsey

## Abstract

Recent studies have identified over one hundred high-confidence (hc) autism spectrum disorder (ASD) genes. Systems biological and functional analyses on smaller subsets of these genes have consistently implicated excitatory neurogenesis. However, the extent to which the broader set of hcASD genes are involved in this process has not been explored systematically nor have the biological pathways underlying this convergence been identified. Here, we leveraged CROP-Seq to repress 87 hcASD genes in a human *in vitro* model of cortical neurogenesis. We identified 17 hcASD genes whose repression significantly alters developmental trajectory and results in a common cellular state characterized by disruptions in proliferation, differentiation, cell cycle, microtubule biology, and RNA-binding proteins (RBPs). We also characterized over 3,000 differentially expressed genes, 286 of which had expression profiles correlated with changes in developmental trajectory. Overall, we uncovered transcriptional disruptions downstream of hcASD gene perturbations, correlated these disruptions with distinct differentiation phenotypes, and reinforced neurogenesis, microtubule biology, and RBPs as convergent points of disruption in ASD.

## Introduction

Autism spectrum disorder (ASD) is an early-onset neurodevelopmental disorder defined by core deficits in social communication and repetitive, stereotyped behaviors^1,2^. While ASD is a genetically and phenotypically heterogeneous disorder, contemporary human genetic studies, focused on rare coding variation, have reliably identified more than a hundred high-confidence ASD (hcASD) genes imparting large effect sizes on risk when disrupted^3,4–6,7–9^. Many of the mutations in these genes are protein-truncating variants (PTVs) and intolerance metrics suggest that hcASD genes tend to be haploinsufficient, suggesting that the primary mechanism of patient-derived variants is loss of function in nature^10^. Thus, methods that block transcription of target genes, such as CRISPR interference (CRISPRi)^11^, are well-suited to model patient-derived variants in hcASD genes. However, the relevant functional consequences of mutations in these genes remain poorly understood.

Evaluating multiple hcASD genes in parallel is critical for identifying convergent phenotypes, which are more likely to reflect core processes disrupted in ASD^8,10^. Accordingly, there is a growing body of *in silico*, *in vitro*, and *in vivo* analyses aimed at identifying points of convergence among ASD-associated genes. Transcriptomic analyses of neurotypical brain tissue spanning multiple developmental stages and anatomical regions have repeatedly demonstrated convergence of ASD genes in developing excitatory neurons of the mid-gestational prefrontal cortex^8,12–16^. Higher-resolution analyses, leveraging transcriptional profiles from laser microdissected tissue or single cells have further implicated mid-gestational excitatory neurons^12,17–20^. Comparative studies using post-mortem brain tissues from ASD individuals and controls also robustly support dysregulation of cortical excitatory neurons in ASD^21–23^. Thus, these descriptive analyses provide strong evidence that neurogenesis of excitatory neurons in the prefrontal cortex is a point of convergent biology underlying ASD. Nonetheless, functional analyses are needed to confirm these observations and to understand the underlying mechanisms. Excitingly, recent advances in high-throughput CRISPR-based functional tools have enabled highly parallelized functional evaluation of a large number of hcASD risk genes in both *in vivo* and *in vitro* model systems.

To date, there have been three major studies conducted *in vivo*, with all three identifying neurogenesis as a point of convergence. Jin *et al.* repressed 35 hcASD genes (selected from Satterstrom *et al.* largely based on expression in human and mouse brain tissue) in mouse embryos using *in vivo* Perturb-Seq, revealing abnormalities in neurogenesis and gliogenesis^24^. Willsey *et al.* leveraged CRISPR to perturb 10 hcASD genes (from Sanders *et al.*^4^) in *Xenopus tropicalis* and identified convergent phenotypes impacting excitatory neurogenesis in the dorsal telencephalon^20^. Mendes *et al*. targeted 10 hcASD genes (selected from Sanders *et al.* based on co-expression in the human brain^12^ as well as SFARI Gene^25^ score) in zebrafish, also highlighting convergence in telencephalic neurogenesis, among other phenotypes^26^.

In addition, several groups have investigated ASD risk genes in parallel using *in vitro* CRISPR-based screens in human neural cell lines^27,28^ or organoids^29,30^. These studies generally focused on smaller sets of manually curated risk genes with varying levels of genetic association, and several filtered for ASD risk genes with a predicted role in transcriptional regulation^27,29^. Additionally, all of these studies evaluated only a single time point and half of them did not generate single cell RNA sequencing data downstream of each genetic perturbation^28,30^. Nonetheless, these studies consistently observed that subsets of ASD risk genes disrupt excitatory^27–29^ and/or inhibitory^29,30^ neurogenesis, in part by delaying or accelerating differentiation^27,28^. They also highlighted several molecular processes as possible points of convergence, though specific processes were not consistently identified across more than one study and the pre-selection of genes reduces the generalizability of these findings. In addition to neural cells, a recent study screened perturbations of 127 manually curated ASD risk genes for disruption of ATP-mediated calcium activity in iPSC-derived astrocytes^31^. Although this study observed functional convergence in that a subset of genes resulted in dysfunctional calcium activity, specific points of molecular convergence were not characterized and the broader relevance of these findings to ASD is unclear.

Overall, despite the highly varied clinical presentation of ASD, the diverse model systems utilized, and the targeting of disparate ASD genes with varying levels of association, these *in silico*, *in vivo* and *in vitro* studies demonstrate a remarkable degree of convergence around (excitatory) neurogenesis. However, the generalizability of these findings to the larger set of hcASD genes, identified with hypothesis-naive exome- and genome-wide approaches, is unclear. Additionally, specific biological pathways underlying these disruptions and relevant to a large number of hcASD genes have not yet been identified.

To begin to address these shortfalls, we leveraged pooled CRISPR interference (CRISPRi) coupled with single cell RNA sequencing (CROP-Seq) in combination with cellular assays to systematically characterize the functional and transcriptional consequences of loss-of-function of 87 hcASD risk genes (chosen from Satterstrom *et al.* solely based on statistical association and without *a priori* assumptions about their biological function) in an *in vitro* model of human cortical neurogenesis (**Figure 1**). Specifically, we generated 2D neural cultures from human induced pluripotent stem cells (iPSCs) and then repressed each of the 87 hcASD risk genes independently via CRISPRi during the early stages of NPC differentiation toward cortical excitatory neurons. We collected three time-points during the most transcriptionally dynamic period of neurogenesis and evaluated the effect of hcASD gene perturbation on differentiation trajectory, cell-state occupancy, proliferation, and transcriptional profile. Overall, we extend previous observations that disrupted excitatory neurogenesis is a key point of functional convergence resulting from ASD-related perturbations^8,20,32–34^, and further implicate tubulin biology^35–44^ and RNA-binding^22,45–52^ as potentially critical molecular processes underlying these phenotypes.

**Figure 1:**
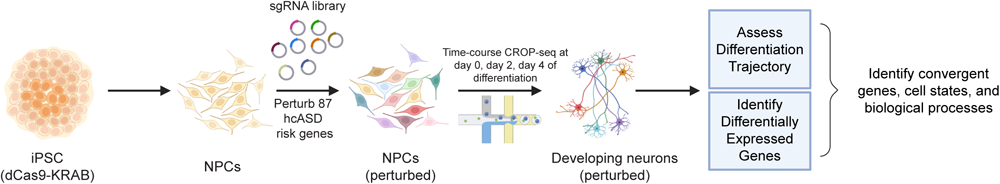
Study Overview. We modeled the impact of hcASD gene perturbations on cortical neurogenesis in vitro, starting with iPSC-derived neural progenitor cells (NPCs). Briefly, we performed a time-course pooled CRISPR interference (CRISPRi) assay coupled with single cell RNA sequencing (CROP-Seq) during the early stages of NPC differentiation to cortical excitatory neurons. We repressed 87 different high confidence ASD risk genes (one per cell) and then utilized single cell RNA sequencing to identify abnormal differentiation trajectories as well as differentially expressed genes. We then conducted follow-up experiments to explore and validate predicted differences in cell cycle, proliferation, and differentiation.

## Results

### Generation of a rapid iPSC-derived 2D neural culture to model human cortical excitatory neurogenesis *in vitro*

Based on the multiple lines of evidence summarized in the introduction, we chose to interrogate the impact of hcASD gene perturbation on excitatory neurogenesis. To model human forebrain cortical neurogenesis *in vitro*, we adapted a small molecule protocol to generate dorsal forebrain-fate neural progenitor cells (NPCs) and cortical deep layer-like excitatory neurons from iPSCs^53^. First, we generated NPCs from iPSCs through dual-SMAD and Wnt inhibition using the small molecules LDN193189, SB431542 and XAV939 (L+SB+X), and maintained NPCs in a proliferative state by supplementing with FGF2 and EGF (**Figure 2A**). We characterized the dorsal forebrain fate of NPCs by examining expression levels of dorsal neural progenitor markers PAX6, SOX2, NES, and forebrain marker FOXG1 with immunocytochemistry (**Figure 2B**). We also confirmed the proliferative capacity of NPCs by quantifying the percentage of Ki67/SOX2 double-positive cells via flow cytometry (**Figure S1A**).

**Figure 2:**
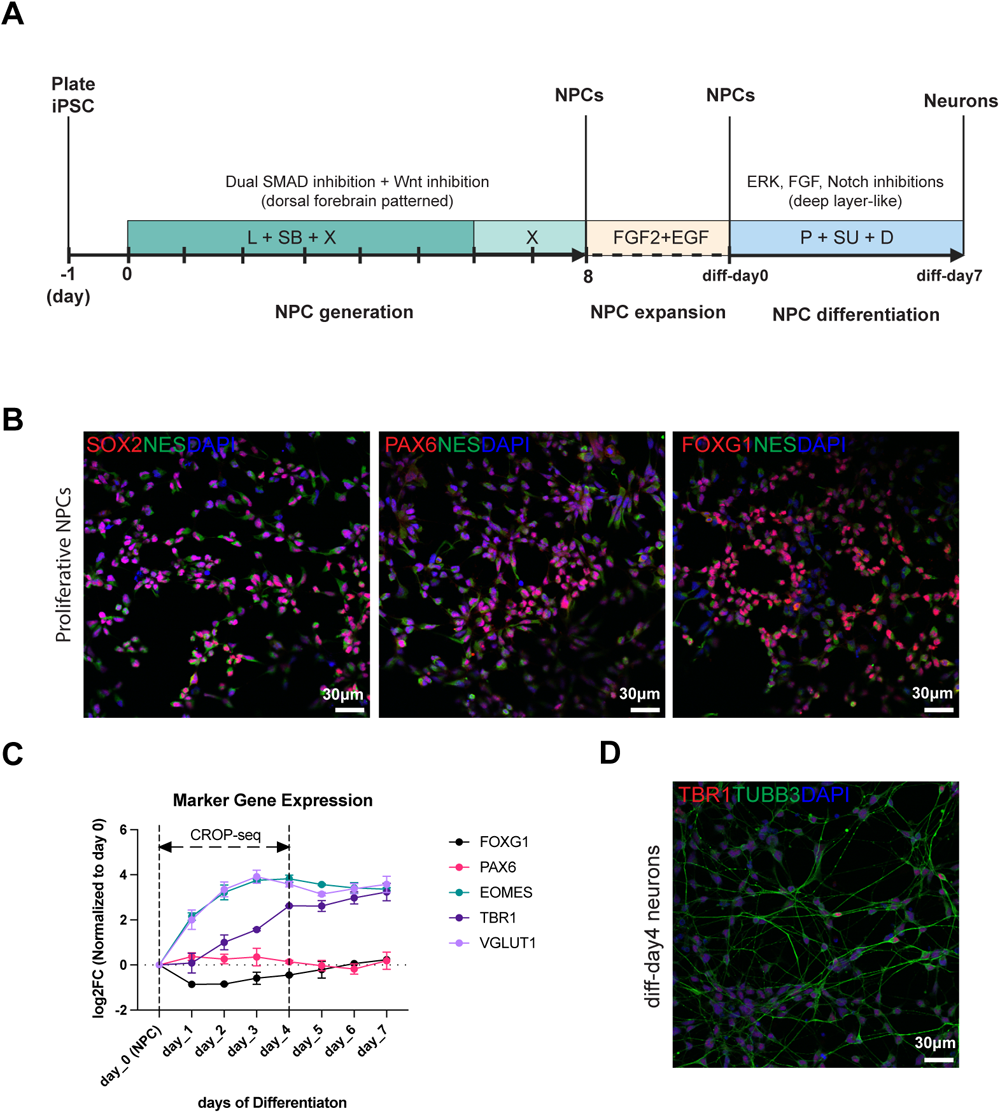
Overview of the iPSC-derived 2D model system of cortical excitatory neurogenesis. (A) Overview of the timelines and protocols we developed for deriving forebrain neural progenitor cells (NPCs) from human induced pluripotent stem cells (iPSCs) as well as for expanding and differentiating these NPCs to immature deep layer excitatory neuron-like cells. We adapted our protocol from a previously developed small molecule based protocol for generating excitatory neurons directly from iPSCs^53^. (B) The NPCs derived by this protocol co-stain for markers of dorsal forebrain NPCs via immunocytochemistry (PAX6, SOX2, FOXG1, and NES). (C) Transcript levels of NPC and excitatory neuronal differentiation markers change dynamically during the first 4 days of differentiation, as measured by qPCR. Markers of more mature excitatory neurons are stably expressed by day 7 of differentiation (e.g. VGLUT1, TBR1). (D) Immature neurons express markers of deep layer cortical neurons by day 4 of differentiation, as assessed by immunocytochemistry against the pan-neuronal marker TUBB3 and the deep-layer marker TBR1. See **Figure S1** for related content.

To generate deep layer-like glutamatergic neurons^8,12^, we cultured NPCs in media containing small molecules PD0325901, SU5402, and DAPT (P+SU+D) to inhibit ERK, FGF and Notch signaling pathways, respectively. We next profiled the transcriptome of cells at day 0, day 7, day 14, and day 22 of differentiation (**Figure S1B, S1C**). We observed that the majority of the 102 hcASD genes identified by Satterstrom *et al.* were within the top 50% of transcribed genes across all four time points (**Figure S1C, 1D**). We also observed that the glutamatergic neuron markers *VGLUT1/SLC17A7*, *TBR1*, and *CTIP2/BCL11B* largely plateaued after 7 days of differentiation (**Figure S1B**), and therefore, we implemented a 7-day protocol to generate immature deep layer-like glutamatergic neurons.

#### Days 2 to 4 of differentiation show the highest dynamic range of differentiation markers

To determine critical time points of cell state transitions within the 7-day differentiation protocol, we measured transcript levels of NPC markers (*PAX6* and *FOXG1*) and differentiation markers (*EOMES/TBR2*, *TBR1*, *VGLUT1*) daily by qPCR (**Figure 2C**). We observed that day 2 was the inflection point for changes in expression level of *VGLUT1*, *EOMES*, and *TBR1* and that by day 4 expression of these genes had plateaued. Moreover, by day 2 of differentiation, we observed a dramatic decrease of occupancy in the G2/M and S phases of the cell cycle, further suggesting that differentiation and some degree of entry into a post-mitotic state may be occurring (**Figure S1E**). Finally, by day 4 of differentiation, we observed, via immunocytochemistry, strong expression of Class III β tubulin (TUBB3) and positive staining for the deep layer neuron marker TBR1 (**Figure 2D**). Taken together, these data suggest that cells are likely undergoing key cell state transitions at day 2 and 4. Thus, we chose these timepoints (as well as day 0 of differentiation) for functional transcriptomic profiling of hcASD gene perturbations.

### Multiplexed knockdowns of high-confidence ASD genes impacts the trajectory of cortical neurogenesis

Using our 2D neuronal differentiation model, we performed a pooled CRISPR interference (CRISPRi) screen of hcASD genes on days 0, 2, and 4 of differentiation of NPCs into cortical excitatory-like neurons, coupled with single cell RNA sequencing (CROP-Seq) ^54,55^. We aimed to repress all 102 hcASD genes identified by Satterstrom *et al*.; however, we eventually excluded 15 hcASD genes from our screen based on low expression in our model system and/or due to low sgRNA knockdown efficiencies after multiple rounds of optimization (see Methods). Thus, the final single guide RNA (sgRNA) library contained sgRNAs targeting 87 of the 102 hcASD genes, with one validated sgRNA per gene) as well as 4 non-targeting control sgRNAs.

#### The 2D model system recapitulates aspects of human cortical excitatory neurogenesis

We recovered a total of 86,551 cells from all three time points across all sgRNAs with an average of 28,850 cells per time point and 951 cells per knockdown. First, we examined target gene repression efficiency in sgRNA+ cells. 73 of the 87 targeted hcASD risk genes had significant on-target repression and/or resulted in one or more differentially expressed genes, whereas the other 14 targeted genes were either not significantly differentially expressed or below the threshold for detection of expression and did not result in any differentially expressed genes **(Figure S2A, S5A)**.

Next, we visualized the clustering of cells in UMAP space and identified 9 clusters using the Leiden algorithm^56^. We then performed an RNA-velocity analysis, using scVelo^54^ (**Figure 3A, S2B**). We also examined the expression pattern of marker genes for cycling (proliferating) and differentiating NPCs (**Figure 3B-C**). Expression of the DNA topoisomerase, *TOP2A*, which is a marker of cycling cells, is consistent with the patterns of RNA-velocity in Leiden clusters 1 and 4 (**Figure 3B, S2B**). Similarly, expression of Doublecortin (*DCX*), which is a microtubule-associated protein with low expression in NPCs and increased expression in developing neurons, is consistent with the patterns of RNA-velocity in Leiden clusters 5-8 (**Figure 3C**). We next performed a velocity pseudotime analysis using CellRank^57^ to associate each cell in the population to a unitless time step between 0 and 1, and observed a strong correlation between pseudotime and expression of *TOP2A* and *DCX* (**Figure 3D, S2C**). Finally, using the marker genes, differential expression of the 9 Leiden clusters, gene set enrichment analysis, and the pseudotime trajectory, we aggregated the cells into 4 macro-clusters encapsulating the major cellular states observed (“cycling”, “intermediate”, “differentiating”; Figure 3E).

**Figure 3:**
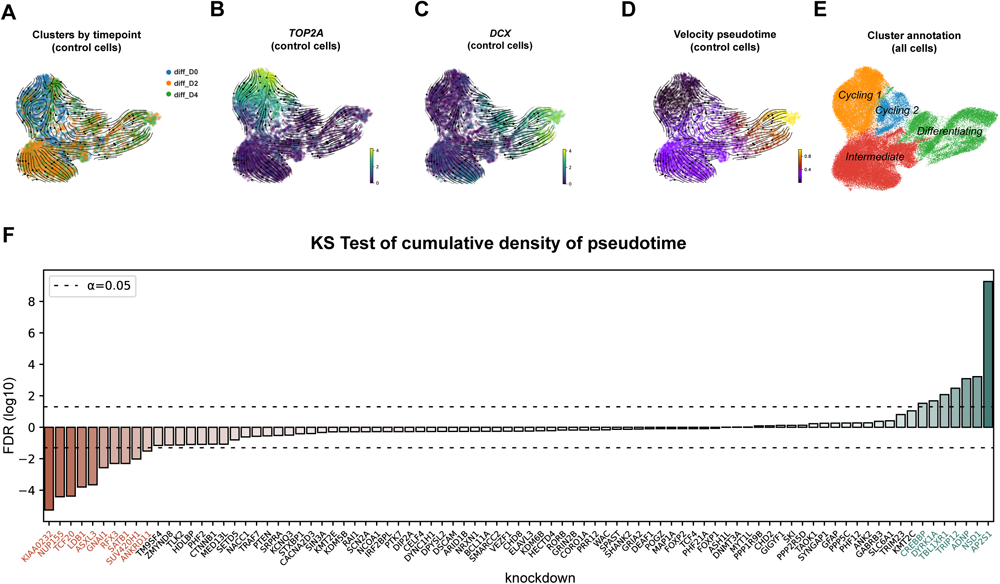
High confidence ASD genes impact the differentiation of neural progenitor cells. (A) UMAP plot of control cells (with non-targeting sgRNAs) from all three timepoints of differentiation, overlaid with RNA-velocity trajectories. (B-C) Expression levels of TOP2A (proliferation marker) and DCX (neural differentiation marker) in control cells, overlaid on the same UMAP as in (A). (D) Velocity pseudotime estimates for control cells, overlaid on the same UMAP as in (A). (E) Annotation of four “macro” cell clusters, identified in part by patterns in A–D and in Figure S2. (F) False discovery rates for hcASD gene knockdown effect on cumulative pseudotime distribution, based on comparison to the cumulative distribution of control cells with non-targeting sgRNAs. Knockdown of 17 genes results in a significant deviation from control cells (so-called ASD pseudotime or ASD_ps_ genes). The 7 genes that result in a “deceleration” of differentiation when knocked down are labeled in green (ASD_dc_) and the 10 genes that result in an “acceleration” of differentiation when knocked down are labeled in red (ASD_ac_). See **Figure S2** for related content.

Finally, we compared the patterns of gene expression observed in our single cell RNA sequencing (scRNA-Seq) data to two sets of scRNA-Seq data generated from human prenatal cortical samples from neurotypical individuals. Comparison with the first dataset, which spans post-conceptional weeks (PCWs) 16 to 24^58^ and captures much of the neurogenesis, demonstrated that our two “cycling” clusters map most closely with “cycling progenitors”, our intermediate cluster maps most closely with “radial glia”, and our differentiating cluster maps most closely with several types of “glutamatergic neurons” (**Figure S2G**). These different cell types map consistently onto our pseudotime trajectory as well. Comparison with the second dataset, which spans PCWs 5.85 to 37^59^, yielded similar results, though the gradient of maturation was less clear when mapping to these data (**Figure S2H**), perhaps because this dataset includes broader stages of neurodevelopment. Importantly, both of these datasets contain cell types from non-excitatory lineages (e.g. interneurons) yet we consistently observe enrichment for excitatory lineage cells only. Taken together, these results suggest that our in vitro iPSC-derived 2D model recapitulates some aspects of excitatory neurogenesis in the human cortex.

#### Pseudotime analysis reveals 17 hcASD genes that impact differentiation trajectory

We next measured the distribution of pseudotime values for cells with the same hcASD gene targeting sgRNA and compared them to the pseudotime distribution of control cells with non-targeting sgRNAs to determine if specific hcASD gene knockdowns affect the differentiation timeline of cells (**Figure 3F**). We identified 17 hcASD knockdowns with significant deviation (FDR < 0.05) from non-targeting controls with respect to their cumulative pseudotime distribution (so-called ASD pseudotime or ASD_ps_ genes). The 17 ASD_ps_ genes have a bimodal pattern of deviation: 7 of the knockdowns have a left-shifted skew in pseudotime, describing a deceleration in differentiation (ASD deceleration or ASD_dc_, labeled in green) and 10 of the knockdowns have a right-shifted skew in pseudotime, describing an acceleration in differentiation (ASD acceleration or ASD_ac_, labeled in red; **Figure 3F, S2F**). To determine if the 17 ASD_ps_ genes simply reflect the most strongly perturbed hcASD genes, we compared the sgRNA knockdown efficiencies for 15 ASD_ps_ genes (*SUV420H1* and *SATB1* do not have detectable expression levels in the scRNA-Seq data but we independently validated their expression and subsequent knockdown in NPCs with qPCR, **Figure S3A**) with 66 of the remaining hcASD knockdowns with detectable expression. We observed that the two gene sets have comparable levels of knockdown as measured by log2FC (**Figure S2D**), suggesting that their identification is based on their biological function rather than extent of perturbation. Additionally, we compared the strength of association with ASD (measured by FDR in Satterstrom *et al*.) for the 17 ASD_ps_ genes and the remaining 70 hcASD and observed that there was no significant difference between the distributions of FDRs for the two gene sets (**Figure S2E**). Similarly, this suggests that the pseudotime analysis pinpointed the 17 ASD_ps_ genes based on their biological function, rather than their effect size in ASD^8^.

The cumulative density plot for the non-targeting control cells has two inflection points, potentially reflecting two differentiation events, which fits well with the pseudotime and cluster maps (**Figures 3D-E, S2F**). Examining the cumulative density of pseudotime for individual knockdowns revealed unique trajectories and how they deviate at specific points from the non-targeting controls (**Figure S2F**). Focusing first on two ASD_dc_ genes: in the case of *AP2S1*, there appears to be a global increase in density in all earlier cell states whereas in the case of *NSD1*, the divergence from non-targeting controls appears right before the second differentiation event. We observed an inverse pattern for two ASD_ac_ genes: *KIAA0232* and *SUV420H1*. *KIAA0232* was consistently accelerated from non-targeting controls throughout differentiation, whereas *SUV420H1* only showed acceleration in pseudotime after the second differentiation event.

### Repression of ASD_ps_ genes disrupts proliferation and differentiation

To better understand the biological correlates underlying the observed accelerations and decelerations in pseudotime, we next conducted several targeted experiments focused on the 17 ASD_ps_ genes. More specifically, we generated separate knockdown cell lines for each of the 17 ASD_ps_ genes as well as for one control cell line with a non-targeting sgRNA (**Figure S3A**) and then assessed cell cycle, proliferation, differentiation, and apoptosis.

#### ASD_ps_ gene knockdowns alter cell cycle

The pseudotime phenotypes of the ASD_ps_ gene knockdowns suggest perturbation of proliferation and/or differentiation, and therefore that cell cycle regulation may be disrupted. Therefore, we asked whether the differential pseudotime phenotypes of the ASD_ps_ genes were associated with cell cycle dysregulation in NPCs and neurons. Durations of G1 and S phase can indicate cell state during neuronal differentiation: proliferative NPCs generally have a shorter G1 phase and a longer S phase than differentiating neurogenic cells^60^. Therefore, we measured cell cycle phase occupancy of bulk cells as an approach to infer durations of cell cycle phases^61^. Specifically, we fluorescently labeled S phase cells as well as DNA content and measured the percentage of cells in each cell cycle phase (G1, S, and G2/M) via flow cytometry to estimate cell cycle phase occupancy (**Figure S1E**). We compared the occupancies in cycling (S+G2/M) and non-cycling (G1) phases between ASD_ps_ knockdown cells lines and the control cell line. For all 17 ASD_ps_ genes, we observed significant deviation in cell cycle phase occupancy in non-differentiating NPCs (proliferating only) and/or differentiating day 2 immature neurons (proliferating and differentiating; **Figures 4A, S3B**).

**Figure 4:**
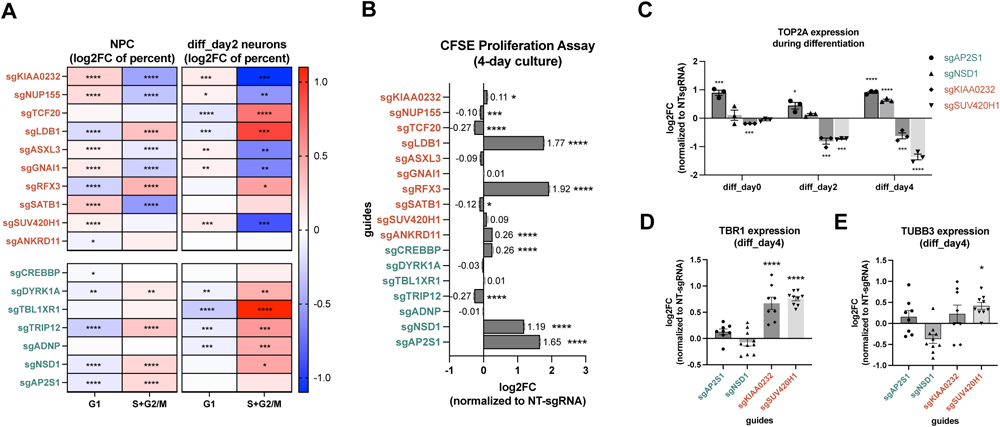
ASD_dc_ and ASD_ac_ gene knockdowns impact cell cycle, proliferation, and differentiation. (A) Heatmap of cell cycle phase occupancies in G1 and S+G2/M of ASD_ps_ cells, expressed as a log2 fold-change over control cells, for non-differentiating NPCs and day 2 Neurons. ASD_ac_ genes are denoted with red text, ASD_dc_ cells are denoted with green text. Red cells denote overrepresentation and blue cells denote underrepresentation. (B) Proliferation analysis using fluorescent CFSE stains comparing ASD_ps_ knockdown NPCs with non-targeting control NPCs, under proliferating conditions. (C) Expression of proliferation marker TOP2A during differentiation for selected ASD_dc/ac_ genes. (D-E) Expression of differentiation markers TBR1 (E) and TUBB3 (F) for selected ASD_dc/ac_ genes on day 4. Asterisks denote level of significance, after applying the Dunnet test to raw p-values: *FDR < 0.05, **FDR < 0.01, ***FDR < 0.001, ****FDR < 0.0001. See **Figure S3** for related content.

In general, repression of ASD_dc_ genes tended to result in a significantly increased percentage of cells in S+G2/M phases in both NPCs and day 2 neurons, suggesting increased proliferation and decreased differentiation, consistent with their observed deceleration in pseudotime. Notably, the onset of this phenotype varied based on the gene. We next separated S and G2/M phases and observed that knockdowns of ASD_dc_ genes mainly increased S phase occupancy and decreased G1 phase occupancy during the NPC stage but exhibited varied cell cycle phase occupancy phenotypes by neuronal differentiation day 2 (**Figure S3B**). However, *CREBBP* knockdown cells are predominantly enriched in G2/M phase and depleted for cells from S phase in NPCs, highlighting a cell cycle disruption distinct from other ASD_dc_ genes.

In contrast, the majority of ASD_ac_ gene knockdowns resulted in an increased percentage of cells in G1 phase and a decreased percentage of cells in S+G2/M phases in both NPCs and day 2 neurons, suggesting decreased proliferation and increased differentiation, consistent with their observed acceleration in pseudotime. However, ASD_ac_ gene knockdowns appear to be much more heterogeneous and temporally dependent in their effects. For example, *LDB1* and *RFX3* knockdowns displayed elevated S+G2/M phase cells and diminished G1 phase cells, paralleling ASD_dc_ genes despite differing pseudotime dynamics **(Figure 3F)**. Interestingly, *ANKRD11* and *TCF20* knockdowns presented unique, temporally specific patterns, underscoring individual gene intricacies.

Overall, these results demonstrate that repression of each of the 17 ASD_ps_ genes disrupt cell cycle phase occupancy. However, while there are general trends of cell cycle phase occupancy resulting from knockdown of ASD_ps_ genes, with ASD_dc_ gene repression correlating with greater occupancy in S+G2/M and ASD_ac_ gene repression correlating with decreased occupancy in S+G2/M, cell cycle phenotype alone is not sufficient to predict differentiation phenotype. This may be due, in part, to the difficulty of separating truly proliferating cells (increased S+G2/M occupancy) from cells stalled during cell division (may also manifest as increased S+G2/M occupancy). Consistent with this possibility, when analyzing S and G2/M phases separately, cells with knockdown of ASD_ac_ genes are much more likely to have a G2/M phenotype than cells with ASD_dc_ gene knockdowns (**Figure S3B**).

#### ASD_ps_ gene knockdowns alter NPC proliferation

To discern whether the pronounced S and/or G2/M phase occupancy in ASD_dc_ knockdowns stems from an increased proliferation rate or cell cycle stalling, we utilized CFSE staining and flow cytometry to assay NPC proliferation under non–differentiating conditions over the course of four days (**Figure 4B**). Notably, *AP2S1*, *NSD1*, and *CREBBP* knockdowns surpassed non-targeting controls in proliferation, suggesting their increased S and/or G2/M phase occupancy corresponds to enhanced proliferation. *TRIP12* knockdown resulted in reduced proliferation, pointing to potential S-phase stalling (**Figure S3B**). The remaining three ASD_dc_ knockdowns did not significantly deviate in proliferation from controls in this 4-day proliferation assay, potentially due to the absence of a clear cell cycle phenotype (*TBL1XR1*, *ADNP*, **Figure 4A**, **S3B**) in NPCs and/or the relatively short time frame of this analysis, which may be insufficient to reveal weak proliferation phenotypes.

In the context of ASD_ac_ knockdowns, those with enhanced G1 phase occupancy coupled with diminished S and/or G2/M phases, such as *SATB1* and *NUP155*, showed decreases in NPC proliferation. Conversely, knockdown of genes like *LDB1*, *ANKRD11*, and *RFX3*, which presented with elevated S and/or G2/M phase occupancy, caused an increase in proliferation in the CFSE assay. Interestingly, despite strong cell cycle phase occupancy phenotypes, we observed no discernible change in short-term proliferation rates for the knockdowns of *SUV420H1, ASXL3,* and *GNAI1*–though this may again be due to the short time frame of this assay. Finally, despite a cell cycle phase occupancy phenotype that would suggest decreased proliferation rate, we observed a slight but significant increase in proliferation rate for the knockdown of *KIAA0232* in NPCs.

Overall, these results suggest that disruption of proliferation is a common feature among ASD_ps_ genes. However, the directionality of this phenotype does not appear to correlate with ASD_dc_ or ASD_ac_ status. In part, this could be due to the fact that we conducted this assay under conditions of proliferation only (i.e. non-differentiating conditions), whereas we identified the pseudotime phenotypes under differentiating conditions, when proliferation and differentiation are entangled. Thus, we next conducted several assays on differentiating cells, focusing on two ASD_dc_ genes (*AP2S1*, *NSD1*) and two ASD_ac_ genes (*KIAA0232*, *SUV420H1*).

#### Expression of proliferative marker TOP2A during differentiation shows a divergence between ASD_ac_ and ASD_dc_ gene knockdowns

We first utilized flow cytometry to estimate the percentage of proliferating (TOP2A+) cells present at day 0, day 2 and day 4 of differentiation. We then calculated the fold-change of proliferating cells in ASD_ac_ and ASD_dc_ gene knockdowns relative to non-targeting controls (**Figure 4C**).

In the ASD_dc_ genes, *AP2S1* knockdown consistently exhibited increased TOP2A+ cell representation throughout differentiation, suggesting a consistent and maintained high proliferation rate. *NSD1* knockdown demonstrated increased TOP2A+ cell percentage starting from differentiation day 4 only, indicating a delayed onset of increased proliferation. Both of these results are consistent with the respective cumulative density of pseudotime plots previously described for these genes (**Figure S2F**).

For the ASD_ac_ genes, *KIAA0232* knockdown persistently revealed a diminished percentage of TOP2A+ cells across all time points of differentiation, indicating attenuated proliferation. *SUV420H1* showcased a notable TOP2A+ cell decline, starting at day 2 and intensifying by day 4, suggesting decreased proliferation during the later time points. Again, these proliferative trajectories coherently align with the differential pseudotime analyses, wherein *KIAA0232* registered an early differentiation increment and *SUV420H1* diverged prominently from controls later in differentiation (**Figure S2F**).

Therefore, in general, the proliferative marker TOP2A appears to separate ASD_dc_ and ASD_ac_ genes with high fidelity. Moreover, its temporal pattern appears to be predictive of differences in pseudotime trajectories, even within ASD_ac_ and ASD_dc_ groups.

#### Immunocytochemistry reveals an increase in neuronal markers for ASD_ac_ genes

Using immunocytochemistry, we assessed neuronal differentiation of the representative ASD_ac_ and ASD_dc_ gene knockdowns by staining for the deep layer neuronal marker, TBR1 (**Figure 4D**), and the pan-neuronal marker, TUBB3 (**Figure 4E**), in day 4 neurons. To quantify the extent of neuronal differentiation, we compared the fluorescence intensity of these markers in cells with ASD_ps_ gene knockdowns to the intensity of these markers in non-targeting controls.

The ASD_ac_ gene knockdowns *KIAA0232* and *SUV420H1* exhibited a pronounced increase in TBR1 fluorescence by day 4 of differentiation and a less pronounced increase in TUBB3. Nonetheless, this suggests that these ASD_ac_ gene knockdowns expedite differentiation relative to non-targeting controls, consistent with their accelerated pseudotime trajectories. Conversely, for both of the ASD_dc_ genes, we did not observe a significant difference in TBR1 or TUBB3 fluorescence, consistent with their decelerated pseudotime trajectories. More specifically, cells with repressed *NSD1* trended towards a decline in TBR1 and TUBB3 fluorescence, implying a potential deceleration in neuronal differentiation. Intriguingly, in the context of the ASD_dc_ gene knockdown *AP2S1*, we discerned a trend towards modest elevation in fluorescence for both TUBB3 and TBR1, though this is most likely due to the enlarged somas of these cells (**Figure S3C**), rather than increased differentiation.

Thus, while the deep layer neuronal marker TBR1 markedly delineates ASD_dc_ and ASD_ac_ genes, the pan-neuronal marker, TUBB3, less clearly separates the two populations of ASD_ps_ genes.

#### hcASD gene knockdowns exhibit temporally distinct apoptotic dynamics during differentiation

To understand how the four ASD_ps_ gene knockdowns impact apoptotic events over time, we quantified the apoptosis marker cleaved caspase 3 (CCP3) using flow cytometry. We then compared the CCP3-positive (CCP3+) cell percentages in gene knockdown samples to non-targeting controls as a means of assessing relative changes in apoptosis (**Figure S3D**).

Our analysis revealed that knockdown of the ASD_ac_ genes *KIAA0232* and *SUV420H1* resulted in an increased percentage of CCP3+ cells at day 0, indicative of early apoptosis. In contrast, at day 0, knockdowns of the ASD_dc_ genes *AP2S1* and *NSD1* either did not change the percentage of CCP3+ cells or significantly decreased it, respectively, suggesting decreased apoptosis. However, this pattern flipped at days 2 and 4. At these timepoints, knockdowns of the ASD_ac_ genes resulted in either a decrease in the percentage of CCP3+ cells (*SUV420H1*) or a return to baseline (*KIAA0232*) whereas knockdowns of the ASD_dc_ genes resulted in either an increase in the percentage of CCP3+ cells (*AP2S1*) or a return to baseline (*NSD1*). Overall therefore, these patterns suggest that decreased early apoptosis may be a component of the ASD_dc_ phenotype, and conversely, that increased early apoptosis may be a feature of the ASD_ac_ phenotype. That being said, more work is needed to understand the generalizability of these findings as well as the unique patterns observed for each gene.

### Cluster overrepresentation reveals convergent cell state changes downstream of hcASD gene knockdowns

To elucidate the overarching cell state changes following hcASD gene knockdown, we assessed whether the distribution of cell counts for each knockdown were enriched or depleted in any of the four macro-clusters (**Figure 3A**). More specifically, within each cluster we compared the distribution for each hcASD gene perturbation to the distribution of non-targeting controls using an internally-developed tool, *cshift*. We present a global overview (collapsing all timepoints) of these shifts in (**Figure 5A**), and time point-specific changes in (**Figure S4A-C**). In all figures, we display all hcASD gene knockdowns with significant enrichments or depletions in one or more timepoints.

**Figure 5:**
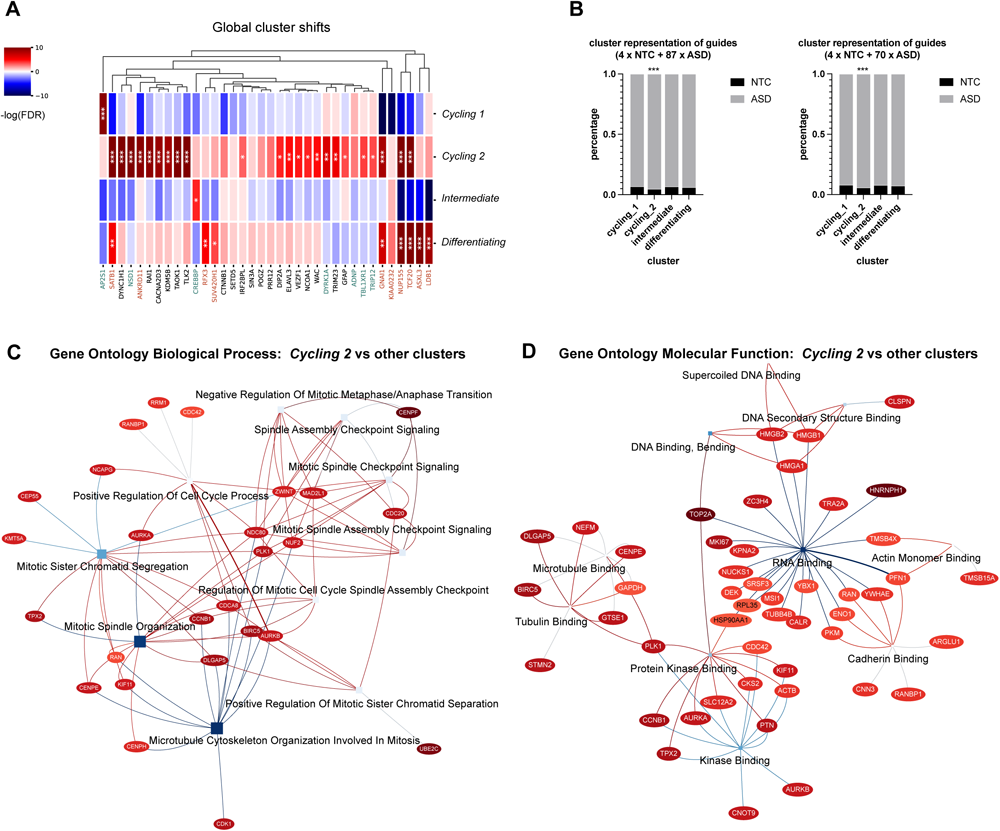
hcASD Gene Knockdowns Converge on a subpopulation of Cycling cells. (A) Cluster representation analysis of hcASD knockdowns reveals contrasting patterns for ASD_dc_ and ASD_ac_ genes, yet a general pattern of overrepresentation in the “Cycling 2” cluster. ASD_ac_ genes are denoted with red text, ASD_dc_ cells are denoted with green text. Red denotes overrepresentation and blue denotes underrepresentation. Significant overrepresentations are indicated by an asterisk. (B) Mean fractional occupancy of all hcASD gene knockdowns across each of the four macro cell clusters, compared to non-targeting controls. “Cycling 2” cells are overrepresented for hcASD gene knockdowns compared to non-targeting controls (left side). This overrepresentation remains even after excluding the 17 ASD_ps_ genes with distinct pseudotime phenotypes (right side). (C-D) Gene set enrichment analysis of genes upregulated in “Cycling 2” cells, using GO Biological Process (C) and Molecular Function annotation categories (D), displayed as IDEA plots where squares denote annotation term names and ellipses are member genes. The top 10 significant terms are shown. Asterisks denote level of significance, after applying the Benjamini-Hochberg procedure to raw p-values: *FDR < 0.05, **FDR < 0.01, ***FDR < 0.001. See **Figure S4** for related content.

#### Differential cluster analysis reveals consistent shifts in hcASD gene knockdowns

Overall, we noted a striking overrepresentation of hcASD gene knockdown cells within the *Cycling 2* cluster (**Figure 5A**). This trend encompassed not only ASD_ps_ gene knockdowns but also numerous other hcASD gene knockdowns. To quantify the significance of this observation and its generalizability to all 87 hcASD genes, we calculated the mean fractional occupancy of all knockdowns across each of the four clusters and conducted a differential representation analysis compared to non-targeting controls. This revealed a prominent overrepresentation of cells with any of the 87 hcASD gene knockdowns in the *Cycling 2* cluster (**Figure 5B**), a result that persisted even after excluding the 17 ASD_ps_ knockdowns (**Figure 5B**). Therefore, this suggests that, in general, disruption of hcASD genes tends to push cells towards the *Cycling 2* “state”. It also raises the possibility that additional hcASD genes may disrupt the trajectory of neuronal differentiation but that we were underpowered to detect their effects in this study.

Despite the convergent phenotype of *Cycling 2* overrepresentation across hcASD genes in general, we still observed a high degree of consistency between shifts in cluster representation and differential pseudotime trajectories (**Figure 5A**). Overall, ASD_dc_ gene knockdown cells tended to be enriched in the *Cycling* populations and depleted in the *Differentiating* population. For example, knockdown of the ASD_dc_ gene, *AP2S1*, resulted in a pronounced overrepresentation of these cells in the *Cycling 1* cluster, matching its slowed differentiation trajectory (**Figure 3F, Figure S2F**) and increased rate of proliferation (**Figure 4A-C**). Likewise, repression of another ASD_dc_ gene, *NSD1*, resulted in a strong overrepresentation of these cells in the *Cycling 2* cluster, consistent not only with its slowed differentiation trajectory and increased rate of proliferation but also with the delayed onset of these phenotypes relative to *AP2S1* (**Figure S2F**). Conversely, ASD_ac_ gene knockdown cells tended to be enriched in the *Differentiating* population. For example, knockdown of the ASD_ac_ genes, *NUP155 and SATB1*, resulted in a clear overrepresentation of these cells in the *Differentiating* cluster, matching their accelerated differentiation trajectory (**Figure 3F**) and decreased number of proliferating cells (**Figure 4C**). That being said, the ASD_ac_ gene knockdowns also tended to be enriched in *Cycling 2* cells, highlighting the complex interplay between proliferation and differentiation as well as what appears to be multifaceted biology of *Cycling 2* cells.

#### Characterization of the *Cycling 2* cluster reveals enrichments in checkpoint signaling, mitotic spindle organization, and RNA binding

To begin to develop hypotheses about the nature of the *Cycling 2* cluster, we identified genes differentially expressed in *Cycling 2* cells, as compared to the cells in all of the other macro-clusters. We then performed gene set enrichment analysis (**Figures 5C-D**).

Our results highlighted significant enrichment in biological processes involved in cell cycle regulation. Specifically, the G2/M checkpoint, the G1/S transition (E2F target genes), and terms related to the mitotic spindle predominated (**Figure 5C, S4E**), consistent with our observations of disrupted cell cycle in cells with perturbed hcASD genes (**Figure 4**). Focusing next on molecular functions, we observed strong enrichment for RNA binding, as well as for microtubule/tubulin, actin, cadherin, and kinase binding, among others (**Figure 5D**).

To better understand putative differences between the cycling clusters, we next compared the *Cycling 2* cluster to the *Cycling 1* cluster. Neuronal differentiation genes like *DCX, STMN2, and STMN4* are upregulated in cells in the *Cycling 2* cluster (**Table S1**). In contrast, when juxtaposed with the *Intermediate* cluster, we observed upregulation of proliferative markers such as *TOP2A, MKI67,* and *CCNB1* (**Table S1**). These results match the velocity pseudotime trajectory across these clusters (**Figure 3D-E**). Taken together, this suggests that the *Cycling 2* cluster may represent a transitional cell state, bridging cycling and intermediate cells. Consistent with this hypothesis, biological processes related to neurogenesis are enriched in the genes upregulated in *Cycling 2* versus *Cycling 1 cells*, and biological processes related to cell cycle, mitosis, and DNA replication are enriched in the genes upregulated in *Cycling 2* and *Intermediate cells* (**Figure S4F-G**).

### ASD_dc_ and ASD_ac_ gene knockdowns have opposing transcriptional signatures

We next leveraged the CROP-Seq data to identify differentially expressed genes (DEGs) by comparing each of the hcASD gene knockdowns to the non-targeting controls. Overall, we identified 3,175 unique DEGs across 73 of the 87 hcASD genes (**Figure S5A**). 1,305 of these DEGs are shared across two or more hcASD genes, 803 are shared across three or more hcASD genes, and 29 are shared across ten or more hcASD genes (**Figure S5B**).

We estimated the effect size of each knockdown by calculating the number of DEGs **(Figure S5A)**. Interestingly, the number of DEGs did not correlate with knockdown efficiency (**Figure S2A**). Along these lines, the 17 ASD_ps_ knockdowns have a much larger number of DEGs compared to the other knockdowns (**Figure S5C**) despite comparable knockdown efficiency (**Figure S2D**), indicating the stronger differential pseudotime phenotypes may be in part due to larger effect sizes of the knockdowns. Indeed, the vast majority of the 3,175 DEGs are present even after narrowing to DEGs of the 17 ASD_ps_ genes only (2,527 of 3,175 or 80%; **Figures 6A, S5D**).

**Figure 6:**
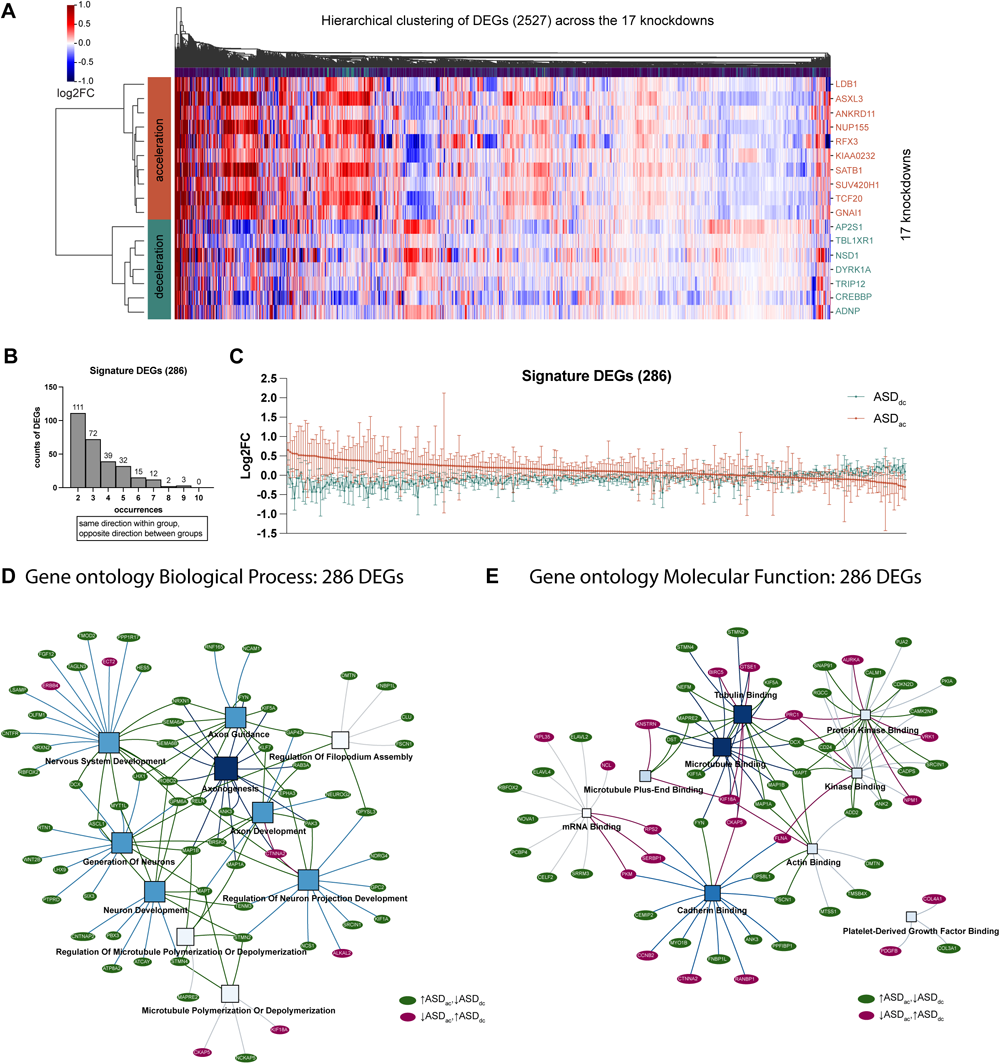
Disruption of ASD_dc_ and ASD_ac_ genes results in opposing transcriptional signatures. (A) Repression of ASD_ac_ and ASD_dc_ genes results in distinct transcriptional profiles. A total of 2,517 differentially expressed genes are present across the 17 ASD_ps_ genes. Hierarchical clustering of the log2 fold change of each of these DEGs for each ASD_ps_ gene knockdown stratifies ASD_ac_ (red bar on left side) and ASD_dc_ genes (green bar on left side). Red cells denote upregulation (positive log2 fold-change) and blue cells denote downregulation (negative log2 fold-change). (B-C) We identified 286 “Signature” genes that have consistently opposite patterns of expression between the ASD_dc_ and ASD_ac_ groups. The plot summarizes the log2 fold-change for each DEG for each ASD_ps_ knockdown, separated by ASD_ac_ (red) versus ASD_dc_ (green), with the within-group median denoted by a horizontal line and the within-group 95% CI denoted by the error bars. Genes are sorted by Log2 fold-change in the ASD_ac_ group. (D-E) Gene set enrichment analysis of the 286 signature genes using Biological Process (D) and Molecular Function (E), visualized as IDEA plots where squares denote annotation term names and ellipses are member genes. The top 10 (or less) significant terms are shown. Genes upregulated in the ASD_ac_ group and downregulated in the ASD_dc_ group are shown in green, whereas genes downregulated in the ASD_ac_ group and upregulated in the ASD_dc_ group are shown in red. See Figure S5 for related content.

We next investigated the 2,517 DEGs underlying the ASD_ps_ gene knockdowns. Strikingly, hierarchical clustering using the log2 fold change of each DEG perfectly stratified the ASD_dc_ and ASD_ac_ genes, with high correlation within groups and inverse correlation between groups (**Figure 6A, S5E**). This suggests that there are convergent transcriptional patterns underlying the differential pseudotime phenotypes and that the majority of transcriptional variation is related to pseudotime.

To investigate this further, we identified 286 “Signature” DEGs that were regulated in opposite directions between the two groups and occurred as significant DEGs in at least two gene knockdowns (**Figure 6B-C**). Interestingly, most of the 286 DEGs are upregulated in the ASD_ac_ group and downregulated in the ASD_dc_ group. Gene set enrichment analysis of these 286 DEGs showed significant enrichment of biological processes related to neurogenesis and axon development (**Figure 6D**). The vast majority of genes in these categories were upregulated in the ASD_ac_ group, consistent with their phenotype of accelerated differentiation. Exploring molecular function, we observed significant enrichment of terms related to mRNA, microtubule/ tubulin, actin, cadherin, and kinase binding (**Figure 6E**). In contrast to the biological process enrichments, genes in the molecular function categories were not consistently up- or down-regulated in the ASD_ac_ group, suggesting more complex dynamics. We also performed gene set enrichment analysis against the MSigDB hallmark gene sets^62^, observing enrichment of terms related to cell cycle, such as “G2/M Checkpoint”, “E2F Targets” (G1/S transition), and “Mitotic Spindle” (**Figure S5F**). Additionally, we observed enrichment for “Hedgehog Signaling”, a pathway not only critical to neural progenitor cell proliferation and differentiation^63–65^ but that has been shown to modulate the effect of hcASD gene disruption during neurogenesis^20^.

Overall therefore, these results are highly consistent with the differential pseudotime and cell cycle phenotypes observed downstream of repression of these genes. They are also highly congruent with enrichments observed in analyses of the *Cycling 2* cluster, which is enriched for hcASD gene perturbations. More specifically, we consistently observe derangements of genes related to cell cycle regulation and the mitotic spindle as well as of genes related to the binding of RNA, tubulin, actin, cadherin, and kinases (**Figures 5A, 6D-E, S4E-F, S5F**).

### Differentially expressed genes are enriched for hcASD risk genes and hcASD protein interactors

We asked whether the transcriptional dysregulation resulting from hcASD gene repression intersects with other ASD-relevant gene sets. We first assessed the overlap between the DEGs identified in this study and newly discovered hcASD genes, based on the hypothesis that hcASD gene knockdowns will converge on ASD-relevant transcriptional changes, and therefore, that the DEGs will be enriched for new hcASD genes. We focused on hcASD genes identified in three new (i.e. published after Satterstrom *et al*., which identified the hcASD genes studied here) exome-wide and/or genome-wide studies of rare variation in ASD^7,9,66^ and assessed overlap with the full set of DEGs (n = 3,175) as well as the subset of “Signature” DEGs (n = 286). We observed that the hcASD genes (FDR < 0.1) identified in the Trost *et al*. and Fu *et al*. datasets are significantly overrepresented within the full gene set (**Figure 7A**). However, hcASD genes from Zhou *et al*. are not significantly enriched, perhaps due to the relatively small number of genes identified in this study (72 versus 255 in Fu *et al*. and 134 in Trost *et al*.) and/or the more moderate effect sizes of the Zhou *et al.* genes^7^. Overall however, the union of the hcASD genes identified across the three studies is also significantly enriched within the full set of DEGs. Notably, after narrowing to the 286 “Signature” genes, we observed significant overrepresentation of hcASD genes from all three studies and a two-fold increase in effect size (**Figure 7A**). Altogether, these results suggest that the DEGs, especially the “Signature” set, are capturing ASD-relevant biology and are predictive of new ASD genes.

**Figure 7:**
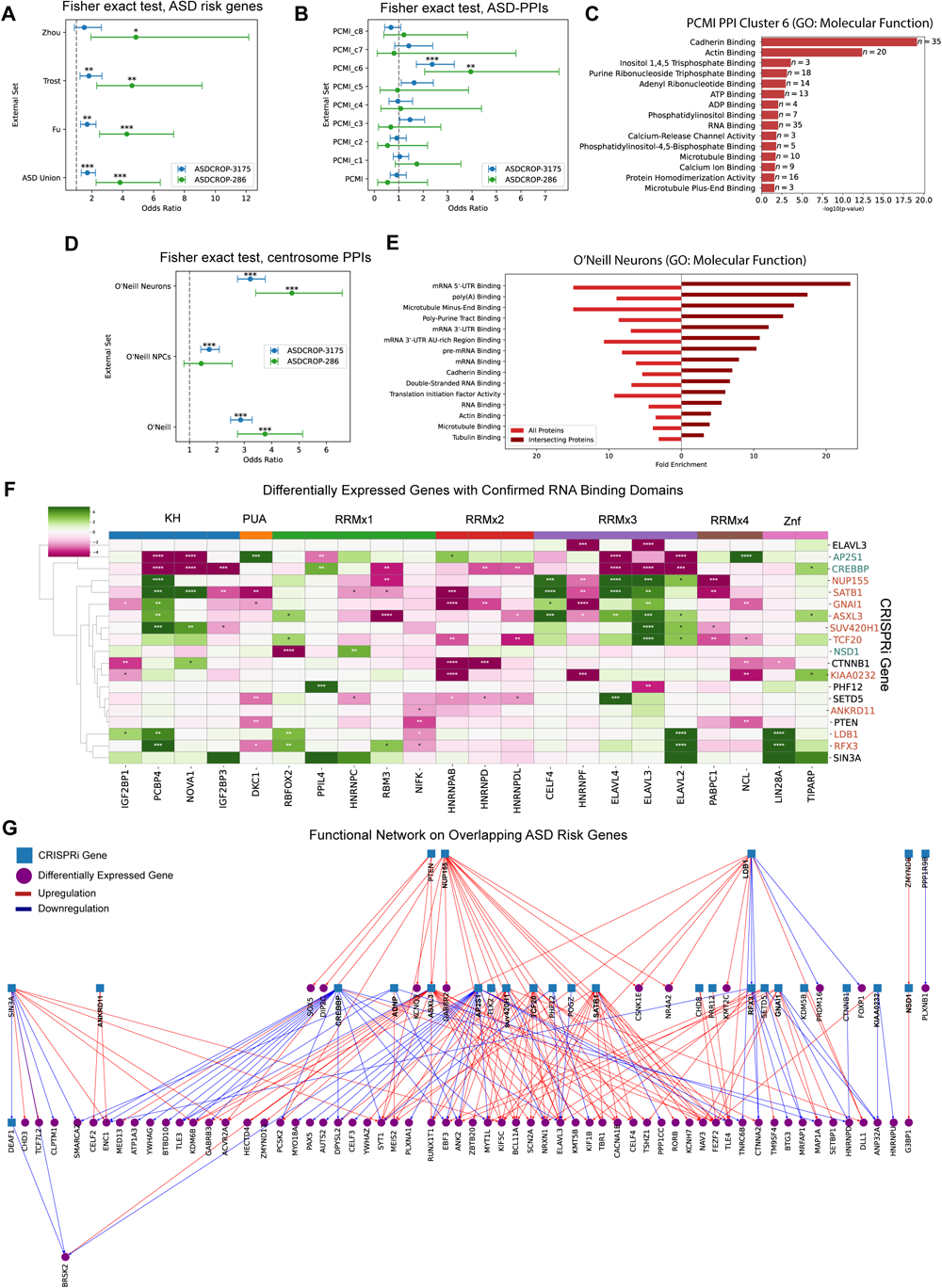
CROP-Seq derived differentially expressed genes overlap with external datasets, suggesting broad relevance and implicating RNA-binding proteins, microtubule-related processes, and centrosomal biology. (A) Differentially expressed genes (DEGs) are enriched for hcASD genes identified in three recent whole-exome and/or whole-genome sequencing analyses of rare variation in ASD^7,9,66^. (B) DEGs are enriched within cluster 6 of an ASD-bait PPI network^67^ (one-sided Fisher’s exact test with Bonferroni correction). (C) Cluster 6 of the ASD-PPI network is enriched for molecular functions related to cadherin, actin, RNA, and microtubule binding. (D) DEGs are enriched within centrosomal interactomes derived from neural progenitor cells and from neurons^44^ (one-sided Fisher’s exact test with Bonferroni correction). (E) The full neuron centrosomal interactome as well as its intersection with the full set of DEGs (n = 3,175) is enriched for molecular functions related to RNA, microtubule/tubulin, cadherin, and actin binding. (F) Many RNA-binding proteins are differentially expressed downstream of hcASD gene perturbations. The heatmap displays experimentally validated RNA-binding proteins^68^ that are differentially expressed within the ASD-CROP-Seq dataset, categorized first by RNA-binding domain and then by the number of occurrences of that domain in the protein (X-axis). The color of the cell corresponds to the “signed” -log10 of the FDR for differential expression downstream of perturbation of a specific hcASD gene (Y-axis), with purple indicating downregulation and green indicating upregulation. ASD_ac_ genes are denoted with red text and ASD_dc_ genes are denoted with green text. (G) A putative functional hierarchy, summarizing the relationship between the hcASD CRISPRi genes from our study (blue squares) and their differentially regulated targets that belong to the union of hcASD genes identified in three recent ASD gene discovery efforts (purple circles)^7,9,66^. Blue arrows indicate downregulation and red arrows indicate upregulation. ASD_ps_ genes are bolded. Asterisks denote level of significance, after applying the Benjamini-Hochberg procedure to raw p-values: *FDR < 0.05, **FDR < 0.01, ***FDR < 0.001, ****FDR < 0.0001. See Figure S6 for related content.

Next, based on the hypothesis that the pathways disrupted downstream of hcASD gene knockdowns will intersect with physical interactors of hcASD proteins, we assessed the overlap between our DEGs and a recently released ASD protein-protein interaction (ASD-PPI) network encompassing 100 hcASD “bait” proteins (85 of which overlap with the hcASD genes studied here), 1,043 “prey” proteins, and 1,881 interactions^67^. We started by dividing this network into 8 clusters based on inter-gene network distance^69^ **(Figure S6A-B, Methods).** We then assessed overlap of the full set of 3,175 DEGs as well as the 286 “Signature” DEGs with the whole ASD-PPI network and with each of the 8 clusters **(Figure 7B)**. We observed significant enrichment of both DEG sets in cluster 6, with the “Signature” gene set again trending towards a larger effect size, indicating that transcriptional changes associated with hcASD gene perturbations are biologically related to a subset of hcASD protein-interactions. Indeed, gene set enrichment analysis of cluster 6 demonstrated enrichment for molecular functions consistent with previous analyses, including cadherin, actin, RNA, and microtubule binding **(Figure 7C)**.

The centrosome is a microtubule organizing center that also functions as a critical hub of RNA modification and interacts with actin^44^. It is also paramount for cell division and dynamically regulated during differentiation of NPCs to neurons (among other cell types). Moreover, recent work has identified significant overrepresentation of hcASD genes in centrosomal PPI networks^36,44^. Thus, given the consistent enrichment of biological processes and molecular functions related to microtubules, RNA-binding, and actin, as well as the role of this organelle in cell division and differentiation, we hypothesized that our DEGs may intersect centrosomal biology. To investigate this, we evaluated the enrichment of the full set of 3,175 DEGs as well as the 286 “Signature” DEGs in centrosomal interactomes recently experimentally derived from NPCs and neurons^44^. Indeed, we observed significant enrichments in both centrosome interactomes, albeit with much stronger signal in neurons **(Figure 7D)**. Hence, we focused on the neuron dataset and conducted gene set enrichment analyses of the full centrosomal interactome as well as of the subset of the interactome overlapping with our 3,175 DEGs (**Figure 7E**). In the full interactome we observed an overrepresentation of molecular functions related to RNA, microtubule/tubulin, cadherin, and actin binding, consistent with the observations of O’Neill *et al*. However, while these terms are also recovered in the gene set enrichment analysis of the intersection between the interactome and our 3,175 DEGs, there was a pronounced shift towards a greater enrichment of RNA-binding related terms (**Figure 7E**). Taken together, these results suggest that the consistent enrichment of terms related to RNA-binding and microtubules observed throughout this study may in fact reflect a convergent point of biology.

Given the particularly strong enrichment of terms related to RNA-binding, we next assessed how individual hcASD gene knockdowns impact known RNA-binding proteins (RBPs). We focused on DEGs with annotated RNA-binding domains and confirmed RNA-binding activity, as curated in the RNA-Binding Protein DataBase^68^ (**Figure 7F**). We observed large dysregulatory events in the RRMx3 and KH families, where genes in these families tended to have opposing patterns of dysregulation between ASD_dc_ and ASD_ac_ genes knockdowns.

We also explored the heterogeneous nuclear ribonucleoprotein (hnRNP) family of RNA binding proteins (RBPs) as 6 of the 22 DEGs (27%) are hnRNP family members (**Figure 7F**) and an additional hnRNP gene, *HNRNPH1*, is significantly upregulated in the *Cycling 2* cluster **(Figures 5D, S4H)**. Moreover, all three of the new (i.e. after Satterstrom *et al*.) exome-wide and/or genome-wide studies of rare variation in ASD identified one or more hnRNPs as hcASD genes^7,9,66^. hcASD gene knockdowns result in widespread dysregulation of several hnRNPs, spanning multiple sub-families (**Figure S6C,D**). Again, some of these genes, like *PCBP4* and *HNRNPC*, were dysregulated across a large number of hcASD gene knockdowns and have opposing patterns in ASD_ac_ versus ASD_dc_ gene knockdowns.

Finally, we visualized the relationship between the hcASD genes perturbed here and hcASD genes identified in three new “hypothesis-naive” sequencing studies of ASD^7,9,66^. More specifically, we narrowed to the 27 hcASD CRISPRi genes with one or more DEGs identified as hcASD genes in Fu *et al*., Trost *et al*., or Zhou *et al* (n = 71). We then generated a functional network, adding directed edges between source genes (hcASD CRISPRi genes) and target genes (DEGs) (**Figure 7G**). This resulting map provides a putative functional hierarchy for the hcASD genes perturbed in our study, with respect to an expanded set of hcASD genes.

For example, many of the hcASD risk genes upregulated *FEZF2,* a transcriptional repressor required for the specification of layer 5 projection neurons in the cerebral cortex^70–73^. Interestingly, repression of *FEZF2* by another hcASD gene, *TBR1*, is critical for restricting the laminar origin of the corticospinal tract to layer 5, thereby ensuring proper development of layer 6 projection neurons^74,75^. Thus, the many hcASD gene perturbations that result in upregulation of *FEZF2*, as well as the perturbations that disrupt *TBR1*, may impact specification of both layer 5 and 6 projection neurons, which fits well with previous work implicating deep layer neurons as a nexus of ASD risk^8,12^. There is also cross-functional control, where different hcASD CRISPRi genes show inverse control over an hcASD gene, like *MYT1L*, a transcription factor involved in neuronal cell differentiation and maturation^76,77^. Remarkably, ASD_ac_ (*SUV420H1*, *TCF20*, *RFX3*) and ASD_dc_ (*CREBBP*, *AP2S1*) genes impacting *MYT1L* have opposing directions of effect. Furthermore, as expression of this gene promotes neuronal differentiation^76,77^, it is intriguing to note that the ASD_dc_ knockdowns result in downregulation of this gene and the ASD_ac_ knockdowns result in upregulation. There are also nested hierarchies within the hcASD gene knockdowns, such as the interaction of *NUP155* and *ASXL3,* where *NUP155* knockdown impacts the expression of *ASXL3*, and as such, they share many downstream targets and have consistent pseudotime and cell cycle phenotypes (**Figure 3F, 4A-B**). Another example of this is two “atypical” ASD_ac_ genes, *LDB1* and *RFX3,* where *LDB1* knockdown impacts *RFX3* expression–though in this case they do not share direct downstream targets despite sharing phenotypes in pseudotime, cell cycle and proliferation **(Figure 3F, 4A-B)**.

### Patient-derived loss-of-function variants in ASD_ps_ genes are associated with brain size phenotypes in human patients

Brain size phenotypes have been associated with ASD in humans, and related phenotypes are often evaluated in *in vitro* and *in vivo* ASD models^20,78–80^. Since ASD_ac_ and ASD_dc_ gene knockdowns led to derangements in multiple cellular processes during NPC differentiation, we hypothesized that patient-derived variants within the 17 ASD_ps_ genes may be associated with altered brain size (macrocephaly or microcephaly) in human patients with pathogenic variants in these genes. Therefore, we queried the DECIPHER database^81^ and tabulated the number of individuals with “open-access” loss-of-function (LoF) sequence variants in the 17 ASD_ps_ genes as well as in a negative control set consisting of the 17 hcASD genes with the lowest pseudotime phenotype (i.e. non-significant and lowest ranked in the K.S test, **Figure 3F**, **Table S4**). The so-called ASD_psNeg_ control genes do not differ from the ASD_ps_ genes in terms of their CRISPRi-mediated knockdown (**Figure S2A**).

Overall, we did not observe a clear relationship between LoF variants in the 17 ASD_ps_ genes and brain size phenotypes, as compared to the 17 hcASD_psNeg_ genes (OR 1.15, p = 0.32, one-sided Fisher’s exact test), though this test is underpowered. However, LoF variants within ASD_dc_ genes alone are associated with a brain size phenotype (OR 1.74, p = 0.019), and this seems to be driven solely by their strong association with microcephaly (OR 2.71, p = 0.00045 for microcephaly versus OR 0.45, p = 0.97 for macrocephaly). In contrast, LoF variants in ASD_ac_ genes alone are not associated with a brain size phenotype (OR 0.69, p = 0.93). That being said, this analysis is hindered by missing data, as 40% (4 of 10) of the ASD_ac_ genes do not have open access data in DECIPHER whereas only 14% (1 of 7) ASD_dc_ genes are missing data. Nonetheless, unlike variants in the ASD_dc_ genes, which are biased towards microcephaly (p = 0.03, two-sided Wilcoxon rank-sum test), variants in the ASD_ac_ genes appear to be relatively evenly distributed across microcephaly and macrocephaly (p = 0.80). This is consistent with our observations that ASD_ac_ gene knockdowns appear to be much more heterogeneous in their impacts on proliferation and differentiation (**Figures 4A-B**, **5A**). Taken together, these results suggest that patient-derived LoF variants in ASD_dc_ genes have more homogenous effects on neurogenesis and tend to result in microcephaly whereas the relationship between patient-derived LoF variants in ASD_ac_ genes and brain size is more complicated (and perhaps obscured by missing data).

## Discussion

Whole-exome and whole-genome sequencing studies of ASD have identified many high-effect high-confidence ASD (hcASD) risk genes^5,66,82^. Yet the cellular and neurodevelopmental consequences of mutations in these genes remain poorly understood, especially with respect to delineating core components of pathology versus unrelated pleiotropic effects^8,10^. Excitingly however, the advent of high throughput, CRISPR-based approaches enables highly parallelized functional studies aiming to link disruptions of these genes to specific molecular processes and phenotypes shared across a large number of hcASD genes, thereby pinpointing convergent points of biology relevant to at least a subset of patients with ASD.

There have been three parallelized *in vivo* screens of a smaller number of hcASD genes (10-35 genes) conducted to date^20,24,26^. All three implicated excitatory neurogenesis, but the specific molecular processes underlying this convergent phenotype were not elaborated. Several more studies have examined ASD risk genes in parallelized *in vitro* screens, similarly identifying neurogenesis-related phenotypes without clear molecular correlates. Additionally, the *in vitro* studies chose genes based on manual curation^27–31^, and sometimes even further restricted to those with similar putative biological functions^27,29^. Although manually-curated gene lists tend to be more comprehensive, they also tend to introduce a degree of bias that is largely avoided by focusing on genes identified by exome- and genome-wide approaches (e.g. hcASD genes)^8,10^. Thus, the generalizability of the *in vitro* findings generated to date is unclear.

Here, we leverage pooled CRISPR interference (CRISPRi) coupled with single cell RNA sequencing (CROP-Seq) to mimic the impact of loss of function variants in 87 hcASD risk genes during the most dynamic period of an *in vitro* model of human cortical neurogenesis, representing the largest screen of hcASD genes conducted to date. Importantly, we assessed multiple timepoints and focused on a large number of hcASD genes, chosen purely based on statistical association, without *a priori* assumptions about their biological function. We also conducted multiple experiments to validate and expand upon the results of the CROP-Seq screen.

Overall, we identify a clear point of convergence around neurogenesis, expanding the generalizability of a large body of work^8,12–24,26–30,32–34^. Specifically, we observe that disruption of hcASD genes in general biases cells towards the *Cycling 2* cluster–a population of cells that may represent a transitional cell state, bridging *Cycling 1* and *Intermediate* cells. Indeed, compared with *Cycling 1* and *Intermediate* cells, we observe differential biology related to neurogenesis, cell cycle, mitosis, and DNA replication. Thus, overrepresentation of hcASD gene knockdowns within this common cell state could indicate differences in proliferation and/or differentiation.

Consistent with this observation, we identified 17 hcASD genes within which disruption leads to clear derangements in cell cycle and differentiation trajectory (ASD pseudotime or ASD_ps_ genes). Ten of these genes resulted in an acceleration of differentiation (ASD_ac_) and 7 of these genes resulted in a deceleration (ASD_dc_), in line with phenotypes observed in other *in vitro* screens of ASD genes^27,28^. Importantly, we identified coherent molecular correlates underlying disruption of the 17 ASD_ps_ genes. All 17 genes have disrupted cell cycles, with generally opposing patterns between ASD_dc_ (increased representation in S+G2/M phases) and ASD_ac_ genes (decreased representation in S+G2/M phases). Interestingly, sorting these 17 ASD_ps_ genes by transcriptional phenotype perfectly stratified the two groups of genes, suggesting that there are convergent transcriptional patterns underlying the differential pseudotime phenotypes. Accordingly, we identified 286 *Signature* genes with opposing transcriptional signatures in the ASD_dc_ and ASD_ac_ gene groups. Notably, when looking at enrichment of biological processes within these genes, terms related to neurogenesis and axon development predominated and these processes were upregulated in the ASD_ac_ group, consistent with their phenotype of accelerated differentiation.

The lack of a clear phenotype in the remainder of hcASD genes could be due to many factors, including insufficient statistical power and/or a requirement for a longer time course to detect weak effects as well as the possibility that these genes are more relevant to different biological processes and/or cell types^30^. Nonetheless, the broader set of differentially expressed genes (DEGs) identified across all hcASD genes significantly overlap with new hcASD risk genes from three recent exome- and genome-wide sequencing studies in ASD^7,9,66^. Similarly, we observed significant overlap between these DEGs and a subnetwork (cluster 6) of a recently generated hcASD protein-protein interaction (PPI) network^67^. Together, these results suggest that the DEGs of hcASD genes converge to some extent as well as capture aspects of ASD biology, suggesting some degree of broad relevance.

Along these lines, in addition to convergence around cell cycle regulation and the mitotic spindle, across orthogonal analyses we recurrently observed enrichment of molecular function terms related to RNA, microtubule/tubulin, actin, cadherin, and kinase binding. More specifically, genes upregulated in *Cycling 2* cells, genes within the 286 *Signature* gene set, and proteins within cluster 6 of the hcASD-PPI network are all enriched for these molecular functions. These consistent enrichments, as well as prior work linking the centrosome with ASD^36,44^, suggested that centrosomal biology may intersect with hcASD gene disruptions during neurogenesis. Indeed, we observed significant overlap between experimentally-derived centrosomal proteins^44^ and both the broader set of DEGs as well as the 286 *Signature* DEGs. This result coherently links our observed neurogenesis phenotypes, as well as the molecular processes putatively underlying them, with the well-characterized role of the centrosome in cell division, cell migration, cilia formation, and neurite outgrowth as well as in organizing RNA-binding/modifying proteins and their targets^44,83–87^. Taken together, these results suggest that the consistent signals observed throughout for microtubule-related biology and RNA-binding proteins are likely interrelated. That being said, it is not clear whether one or both are driving the phenotypes we have observed here. Thus, more work is needed to disentangle the relevance of microtubule biology versus RNA-binding proteins.

Furthermore, these are relatively broad functional categories. For instance, within microtubule-related biology, many of the signals observed throughout this paper suggest that the proliferation and differentiation phenotypes could be due to disruption of the mitotic spindle, especially as a subset of hcASD genes have recently been shown to localize to the mitotic spindle^36^. That being said, our gene set enrichment analysis of the 286 *Signature* genes also highlighted Sonic hedgehog signaling, which intersects hcASD gene disruption during neurogenesis^20^ and is localized to primary cilia^88,89^, a microtubule-based organelle essential for NPC proliferation and differentiation^63–65^. As the content and function of the centrosome changes during neurodevelopment^44,90–92^, perhaps the much stronger overlap between the DEGs identified here and the neuron centrosomal proteins offers some insight. Even so, careful experimentation will be required to parse the relative contributions of microtubule-related structures to hcASD-associated disruptions in proliferation and differentiation during neurogenesis.

Finally, despite the fact that individuals with ASD demonstrate extremely heterogeneous clinical presentations, we observed a strong relationship between ASD_dc_ genes and microcephaly. However, much more work is needed to understand how ASD_ps_ phenotypes *in vitro* relate to neurodevelopmental phenotypes in human patients. Nonetheless, this raises some hope that *in vitro* screens of ASD genes, particularly those chosen in a hypothesis-naive manner, have the potential to generate insights with meaningful clinical correlates, and therefore, that they may eventually help to stratify patients and/or inform the development of novel treatment strategies and targets.

## Acknowledgements

This research was funded by grants from the National Institutes of Health to M.K. and A.J.W (U01MH115747), A.J.W. (U01MH116487), and B.W. (R25MH060482). This study was also supported by the Weill Institute for Neurosciences (Startup Funding to A.J.W.), the Overlook International Foundation (to A.J.W.), and the Sorensen Foundation Career Award in Child & Adolescent Psychiatry (to B.W.). H.R.W. is a Chan Zuckerberg Biohub - San Francisco Investigator. The authors would like to thank all members of the Psychiatric Cell Map Initiative (PCMI; U01MH115747), the Willsey Labs, and the Kampmann Lab for their invaluable discussions and support.

## Author contributions

Conceptualization: N.S., N.Teys., R.T., M.K., A.J.W.; Methodology: N.S., N.Teys., R.T., M.K., A.J.W.; Software: N.Teys.; Formal Analysis: N.S., N.Teys., A.E.; Investigation: N.S., S.D., M.S., Y.Z., N.Teer., R.T.; Resources: M.K., A.J.W.; Data Curation: N.Teys., A.E.; Writing – Original Draft: N.S., N.Teys.; Writing - Review & Editing: N.S., N.Teys., B.W., S.D., M.S., Y.Z., A.E., N.Teer., H.W., H.G., R.T., M.K., A.J.W.; Visualization: N.S., N.Teys.; Supervision: H.G., R.T., M.K., A.J.W.; Project Administration: M.K., A.J.W.; Funding Acquisition: B.W., M.K., A.J.W.;

## Materials and Methods

### Cell Culture

#### iPSCs

We obtained the iPSC line engineered with dCas9-KRAB machinery from Allen Cell Collection (Cell line ID: AICS-0090 cl.391). The iPSCs were maintained in mTeSR medium (STEMCELL Technologies, Cat#100-0276) on Matrigel (Fisher Scientific, Cat#08774552) coated tissue culture plates.

#### Generating Neural Progenitor Cells (NPCs) from iPSCs

Dorsal forebrain fate neural progenitor cells were generated from iPSCs through dual SMAD inhibition and WNT inhibition using small molecules (adapted from Qi *et al*., 2017^53^). At day -1, iPSCs were plated on Matrigel coated plates at ∼400k cells/cm^2^ in mTeSR plus medium (STEMCELL Technologies, Cat#100-0276). From day 0 to day 3, cells were fed with KSR medium (15% Knockout Serum Replacement in Knockout DMEM, 1xGlutaMAX, 1xMEM-NEAA, 0.1mM BME) containing small molecules 250nM LDN193189 (LDN) (Tocris, Cat No. 6053), 10uM SB431542 (SB) (Tocris, Cat#1614) and 5uM XAV939 (XAV) (Tocris, Cat#3748). At day 4, cells were cultured in ⅔ KSR + ⅓ N2 (DMEM/F12, 1x N2, 1x B27 -Vitamin A, 1x GlutaMAX, 1x MEM-NEAA) + LDN/SB/XAV, and at day 5 medium switched to ⅓ KSR + ⅔ N2 + LDN/SB/XAV. At day 6, cells were passaged with EDTA at 1:2 and cultured on Matrigel coated plates in NPC medium containing DMEM/F12, 1xN2, 1xB27 -Vitamin A, 1x GlutaMAX, 1x MEM-NEAA, 10ng/ml FGF2 (Peprotech, Cat#100-18B) and 10ng/ml EGF (Fisher Scientific, Cat# 236EG200) supplemented with 5uM XAV. At day 7, cells were fed with the same medium as day 6. At day 8, cells were passaged at 1:3 using Accutase (STEMCELL Technologies, Cat#07920) and cultured in NPC medium on Matrigel coated plates onwards.

#### Generating neurons from NPCs

Deep layer-like cortical excitatory neurons were generated from NPCs through inhibitions of ERK, FGF and Notch signaling pathways using small molecules (adapted from Qi *et al.*, 2017^53^). Specifically, NPCs were plated onto Poly-L-Ornithine (PLO)(MilliporeSigma, Cat#27378-49-0),laminin (Fisher Scientific, Cat#23017015), and fibronectin (MilliporeSigma, Cat#F1141-1MG) coated plates at ∼270k cells/cm^2^ in NPC medium at day -1. At day 0, cells were fed with neural differentiation medium (NDM: neurobasal medium with 1 x B27 no vitamin A, 1 x N2, 1 x GlutaMAX, 1 x MEM-NEAA, 20ng/ml BDNF, 0.5mM dibutyryl cAMP, 0.2mM ascorbic acid) containing small molecules 1uM PD0325901 (P)(Tocris, Cat#4192), 5uM SU5402 (SU)(MilliporeSigma, Cat#SML0443), 10uM DAPT (D)(Tocris, Cat#2634). Media was refreshed every three days. For long term differentiation, small molecules were withdrawn at day 7 and onward.

#### Establishing CRISPRi NPCs for validation experiments

##### Single guide RNA (sgRNA) design and cloning

Top and bottom oligos of non-targeting sgRNAs and ASD risk gene-targeting sgRNAs were designed using a CRISPRi guide RNA design tool^93^ (https://github.com/mhorlbeck/CRISPRiaDesign) developed by Martin Kampmann lab, and were ordered from IDT. Top and bottom oligos were annealed and the annealed products were inserted into a linearized lentiviral vector pMK1334 (Addgene, Cat#127965) through ligation. The ligation products were transformed into DH5 Alpha competent cells. Cells with the desired plasmid were selected using ampicillin and amplified. DNAs of lentiviral vectors carrying sgRNAs were purified from cells using QIAprep Spin Miniprep kit (Qiagen, Cat#27106) and were Sanger sequenced for sequence validation.

##### Lentiviral production and NPC transduction

To establish individual hcASD gene knock-down NPC lines for targeted experiments, NPCs with CRISPRi machinery were transduced with a lentiviral vector carrying a targeting sgRNA. Specifically, lentiviruses were produced in Lenti-X 293T cells (Takara Bio, Cat#632180) and lentivirus-containing supernatants were concentrated using Lenti-X concentrator (Takara Bio, Cat#631231). NPCs were transduced and selected using 4ug/ml puromycin (Fisher Scientific, Cat#5015328) to obtain greater than 90% transduced cells in culture. The transduction efficiency was evaluated by measuring BFP expression in the sgRNA lentiviral vector by flow cytometry. Knock down efficiency of the sgRNAs for the 17 ASD_ps_ genes were validated by qPCR in NPCs under non-differentiating conditions, i.e. conditions meant to maintain proliferation (**Figure 2A**; see also next section). Specifically, we extracted total RNA from NPCs transduced with non-targeting control sgRNA and hcASD targeting sgRNA as described above. Samples were collected in triplicates for each condition. We then converted the total RNA into cDNA through reverse transcription. We used cDNA to measure the target gene expression using the double delta Ct method and calculated the log2FC of a target gene in hcASD gene knocked down cells compared to the control cells.

#### Validation of expression and knockdown of the 17 ASD_ps_ genes in NPCs

Previously, when we assessed knockdown efficiency of the 17 ASD_ps_ knockdowns using data from the CROP-seq experiment, expression of *SUV420H1* and *SATB1* did not pass the detection threshold, and knockdowns of *ASXL3, NUP155, TCF20* and *DYRK1A* were not statistically significant **(Figures S2A, S5A)**. However, the large number of DEGs for these ASD_ps_ genes suggested that we achieved a meaningful degree of repression. Therefore, we utilized qPCR to perform an orthogonal check of knockdown efficiency, and confirmed detectable expressions of all 17 genes in NPCs and significant knockdown of *SUV420H1*, *SATB1*, *TCF20*, and *DYRK1A* as well as borderline significant knockdown of *NUP155* (**Figure S3A**). sgRNAs against the other 11 previously confirmed ASD_ps_ genes also validated.

### CROP-seq

#### CROP-seq sgRNA library construction

First, we designed non-targeting/control and hcASD gene targeting single guide RNAs (sgRNAs) using a pipeline that the Kampmann lab developed^93^(https://github.com/mhorlbeck/CRISPRiaDesign). An average of two sgRNAs were designed for each target gene. We then validated the majority of the sgRNAs either through establishing individual knockdown cell lines and measuring the target gene expression by qPCR, or through a pilot CROP-seq experiment in NPCs. We preferably selected sgRNAs that have knockdown efficiency between 30% and 70%. The hcASD gene sgRNA sequences are listed in **Table S5**. For the CROP-seq experiment presented in this study, we included one validated sgRNA per gene. We conducted pooled cloning to construct a CROP-seq sgRNA library that allows constitutive expression of sgRNAs in cells. Specifically, we digested the pMK1334 lentiviral vector with BstXI + BlpI restriction enzymes. Linearized pMK1334 vector was purified using NucleoSpin Gel and PCR clean-up kit (Takara Bio, Cat#740609.50). Top and bottom oligos of non-targeting and targeting sgRNAs were annealed. Annealed sgRNAs were pooled using the desired amount. The pooled annealed oligos were then diluted and inserted into linearized pMK1334 lentiviral vectors. A test transformation was performed in DH5 Alpha competent cells to assess the ligation efficiency, followed by a large-scale transformation in Stellar Competent cells for final library amplification and purification. Before transducing cells, we measured the proportion of each sgRNA in the library through sgRNA enrichment PCR and high throughput sequencing to check the sgRNA representations in the library. For sgRNAs with low counts, we spiked them into the existing library to ensure no dropouts of sgRNAs.

#### Cell preparation

We produced a pool of lentiviruses containing the CROP-seq sgRNA library and transduced NPC with relatively low titer of lentiviruses (10-20% transduction efficiency) to minimize multiple infections in which a cell receives more than one sgRNA. NPC cultures after puromycin selection were used for the CROP-seq experiment. At day 0, day 2 and day 4 of differentiation, cells were harvested using papain (Worthington, Cat#9001-73-4). Cells were prepared according to 10x Genomics general sample preparation protocol.

#### Single cell RNA-seq library preparation

At each time point, ∼90k cells were loaded into 3 wells (∼30k cells per well) of the Chromium Next GEM Chip G for GEM preparation. GEMs of each time point were stored in -20°C. Single cell library preps of all 9 samples from the three time points were conducted together according to the protocol provided for Chromium Next GEM Single Cell 3’ Reagent Kit v3.1 (10x Genomics, Cat#PN-1000121). Concentrations of libraries were measured using Agilent High Sensitivity DNA kit (Agilent Technologies, Cat#5067-4626) on Bioanalyzer.

#### sgRNA enrichment

Briefly, three PCR reactions were performed followed by 1x SPRIselect beads (Fisher Scientific, Cat#NC0406407) cleanup after each PCR reaction. In PCR1, 15ng full-length cDNA from single cell RNA-seq library prep per sample was used as a template. In PCR2, 10ng post-cleanup PCR1 product was used as a template. In PCR3, 10ng post-cleanup PCR2 product was used as a template to add sample indices. All PCR reactions were conducted using KAPA Hotstart HiFi ReadMix (VWR, Cat#103568-584) with annealing temperature at 62°C (15 sec) and extension at 72°C (15 sec) for 18 cycles (PCR1) or 15 cycles (PCR2 and PCR3). Concentrations of post-PCR3 products were measured using Qubit dsDNA HS assay kit (Thermo Scientific, Cat#Q32854).

#### Sequencing

Samples of single cell RNA-seq libraries and enriched sgRNAs were pooled together at 5nM for sequencing on S2 and S4 flow cells of Illumina NovaSeq. Approximately 1 billion reads were generated per sample.

#### Sequence Alignment

A kallisto index was generated from the human transcriptome using the ENSEMBL cDNA reference (GRcH38). Introns were annotated using the GRCh38 version 105 genome annotation. The default k-mer size (k=31) was used in generation of the index. The 10X sequences were then pseudo-aligned to this index using kallisto-bustools ^94^. The ‘lamanno’ workflow was specified, and the reads were mapped to both spliced and unspliced transcripts. After pseudo-alignment, barcodes were then corrected and filtered against the 10X-v3 cell barcode whitelist. Total gene level counts were calculated by aggregating both spliced/unspliced counts.

#### Knockdown Demultiplexing

A kallisto index was generated for the sgRNA library with a k-mer size of 15. The sgRNA enrichment PCR sequences were then pseudo-aligned to this index using kallisto-bustools ^94^. After pseudo-alignment, barcodes were corrected and filtered against the 10X-v3 cell barcode whitelist. Cells were then assigned to guides using geomux (https://github.com/noamteyssier/geomux), which performs a hypergeometric test for each cell on its observed guide counts, then calculates a log2-odds ratio between the highest counts. Cells were assigned to their majority guide if their Benjamini-Hochberg corrected P-value was below 0.05, the log-odds ratio was above 1, and the total number of UMIs were greater than 5.

#### Single-Cell Analysis

##### Preprocessing

Cells were merged for the 10X sequence and the enrichment PCR by matching on cell-barcode and GEM library. Cells were filtered to only include those with a minimum count of 1000. 3000 of the most highly variable genes were selected with a minimum shared count of 20. Cells were then normalized to a target sum of 1000 and log transformed. Principal component analysis was then performed using these highly variable genes. These components were then batch corrected using the time point of the cells using Harmony ^95^. Nearest neighbors were then calculated using the top 30 principal components, selecting the 30 nearest neighbors. Clustering was performed using the Leiden algorithm, setting the resolution to 0.5 ^56^. A cluster of fibroblast-like cells were then removed from the analysis (filtered by their Leiden cluster ID). All single-cell filtering, transformation, and dimensionality reduction was performed using scanpy ^96^.

##### RNA-Velocity

RNA-velocity analysis was performed using scvelo^54^. Each cell’s nearest neighbors were calculated using the top 30 principal components and used their closest 30 neighbors. The moments were then computed using the top 30 principal components and the closest 30 neighbors. The dynamics of the system were then recovered, and the velocity analysis was performed using the stochastic mode in scvelo. The velocity graph was then computed.

##### Velocity Pseudotime

The velocity pseudotime analysis was performed by first identifying probable initial and terminal states. These states, and their corresponding cells, were determined using CellRank ^57^ - setting the cluster key to the Leiden cluster ID. The highest likelihood initial cell and the highest likelihood terminal cell were then used as the root and end key respectively for the velocity pseudotime function within scvelo.

##### Cluster mapping using scRNA-seq data from primary human cortical tissues

To see how the transcriptomic profiles of the cells in the CROP-Seq we calculated a cluster score for each of our cells against the transcriptomes of clusters identified from a dataset of single cell RNA sequencing of human prenatal cortical samples of neurotypical individuals spanning post-conceptional weeks 16-24 and a dataset of single-cell sRNA sequencing spanning post-conceptional weeks 5.85-37. We calculated a cluster score by identifying the marker genes for each of the clusters identified in these datasets and then measuring the average expression those genes subtracted with the average expression of a reference set of genes (the reference set being a set of randomly selected from all genes and stratified by expression bins). This was done using the scanpy ‘scanpy.tl.score_genes’ function and is a reproduction of the approach taken by Satija et al. 2015^96,97^.

##### Differential Pseudotime

Every cell in the dataset has an associated sgRNA knockdown and pseudotime, so a distribution of pseudotime values was created by grouping all cells by their associated sgRNA. The distribution of each knockdown sgRNA was then compared to the distribution of the non-targeting control sgRNAs via a Kolmogorov-Smirnoff test (ks-test). The associated p-values for the KS-tests were then corrected for multiple hypothesis testing using the Benjamini-Hochberg step-up correction. Significant tests were selected by selecting sgRNAs with an adjusted p-value lower than 0.05.

##### Differential Expression Analysis

Differential expression analysis was performed using different filtering criteria than the single-cell analysis mentioned above. Cells were filtered using a minimum count of 3000 and genes were filtered to those that had a minimum shared count of 10. The fibroblast-like cells were then removed (identified using the previous methodology). These cells were then grouped by knockdown, timepoint, and GEM-library using ADPBulk (https://github.com/noamteyssier/adpbulk). Differential expression tests were performed using DESeq2^98^ comparing each sgRNA knockdown against the non-targeting control sgRNA knockdowns.

##### Cluster overrepresentation analysis

We calculated Leiden cluster overrepresentation for each guide using a chi-square test for each guide/cluster in the dataset. Specifically, a chi-square test is performed for each knockdown group and Leiden cluster between each group distribution and each non-targeting control distribution. The p-values from this are then aggregated over the non-targeting controls using a geometric mean. Finally these p-values are adjusted for multiple hypothesis testing using a Benjamini Hochberg correction. We performed this analysis using the open-source python module cshift (https://github.com/noamteyssier/cshift)

##### Knockdown Clustering Analysis and Gene Set Enrichment

A subset list of knockdowns was created using those that were significant from differential pseudotime analysis. A subset list of differentially expressed genes was then created using the union of genes differentially expressed in each of the knockdowns. Pairwise spearman correlation was then computed for all knockdown pairs using their log2 fold changes for the joint set of differentially expressed genes. The dendrograms were computed using a complete linkage, cosine metric, hierarchical clustering method on the pairwise correlation matrix. Gene set enrichment analysis was performed using Enrichr^99^ and visualized using IDEA (https://github.com/noamteyssier/idea).

##### Comparison to external datasets

We compared differentially expressed genes from the ASD-CROPSeq dataset to external datasets using a one-sided (greater) Fisher’s exact test using all the tested differentially expressed genes as a background (n = 17,542). Odds ratios are calculated from the contingency tables and confidence intervals bounds (0.95) are calculated via the odds_ratio and confidence_interval functions in Scipy. ASD-CROPSeq differentially expressed genes were selected using an FDR ≤ 0.05 (n = 3,175). The n = 286 set was calculated by selecting all genes that were significantly differentially expressed in the same direction within either the ASD_dc_/ASD_ac_ group but inversely expressed in the ASD_dc_/acASD_ac_ group (i.e. upregulated in one and downregulated in the other or vice versa). hcASD genes from Fu *et al., Zhou et al., and Trost et al.* were selected based on a cutoff of FDR ≤ 0.1. O’Neill *et al.* proteins were selected based on a threshold of p value ≤ 0.05.

### CFSE proliferation assay

NPCs were stained with CFSE dye from the CellTrace CFSE proliferation kit (ThermoFisher Scientific, Cat# C34554) according to the manufacturer’s protocol. Stained NPCs were plated in 3 separate wells per line at day 0 and grown for 4 days. From day 0 to day 4, cells were collected daily and stained with live/dead fluorescent dye before being fixed with 4% PFA. The fluorescent intensity of CFSE was measured for all samples by flow cytometry. The geometric mean of the CFSE peak was taken to calculate relative proliferation rate for each sample.

### Flow cytometry

#### EdU labeling for cell cycle analysis

Cells were incubated in NPC media containing 10uM EdU for 1hr. Labeled cells were detached with either Accutase for NPCs or Papain for neurons. EdU was stained using Click-iT™ Plus EdU Alexa Fluor™ 488 Flow Cytometry Assay Kit (Thermofisher Scientific, Cat#C10633).

#### Intracellular stain

For intracellular staining of transcription factors during NPC characterization, cells were fixed and permeabilized with eBioscience FOXP3/Transcription factor staining buffer set (Thermofisher Scientific, Cat#00-5523-00).

Cells were analyzed using BD LSR-Fortessa flow cytometer and data was analyzed using FlowJo software. A list of primary antibodies were provided in **Table S5**.

### Immunocytochemistry

Cells grown on coverslips were fixed with 4% paraformaldehyde for 10 min at room temperature (RT) and permeabilized with 1 x PBS solution containing 0.2% Triton X-100 for 10min. Cells were blocked in 1x PBS + 0.1% Triton X-100 solution containing 5% normal donkey serum for 30 min at RT. Primary antibodies were diluted in the blocking buffer and incubated with cells at 4°C overnight. Fluorophore conjugated secondary antibodies were diluted in blocking buffer and incubated with cells at RT for 1hr. DAPI was used for nuclear counterstains. Stained cells were mounted using Prolong™ Glass Antifade Mountant (Thermofisher Scientific, Cat#P36982) and stored at 4°C. Cells were imaged using Leica SP8 laser scanning confocal microscope and raw images were processed and analyzed using FIJI and CellProfiler. A list of primary antibodies were provided in **Table S5**.

### Quantitative PCR

Total RNAs were extracted from cells using RNeasy Mini Kit (Qiagen, Cat#74106) and were converted into cDNA using SUPERSCRIPT IV VILO MASTERMIX (Thermofisher Scientific, Cat#11756050). Quantitative real-time PCR was performed using Applied Biosystems PowerUp SYBR Green Master Mix (Fisher Scientific, Cat#A25742) on QuantStudio6 real-time PCR system. A list of primers were provided in **Table S5**.

### Bulk transcriptome profiling and analysis

Duplicate total RNA samples were collected for each condition. cDNA converted from ∼10 ng total RNA per sample was diluted in water to make a final volume of 15ul. cDNA samples were then added into wells of IonCode 96 Well PCR plate. Ion AmpliSeq transcriptome human gene expression panel (Thermofisher Scientific, Cat#A31446) was added to the Reagents cartridge. We used the Ion Chef System for automated library preparations with the Ion 540 kit (Thermofisher Scientific, Cat#A30011). Pooled barcoded libraries were subsequently loaded into the Ion GeneStudio S5 sequencer for high throughput sequencing to obtain 60-80 million reads per run.

### Diffusion State Distance (DSD) network proximity

An ASD protein-protein interaction network (ASD-PPI) was previously generated by individually overexpressing 100 Strep-tagged hcASD genes (as “bait” proteins) in HEK293T cells in biological triplicates and identifying “interactor” proteins by affinity purification followed by mass spectrometry 48 hours after transfection^67^. We defined 8 subclusters within ASD-PPI by defining sets of hcASD bait that were more closely connected in protein interaction space. Clustering of the baits was based on network distances calculated by combining the PPI network, all edges weight = 1, with a subnetwork of STRING, and computing the Diffusion State Distance (DSD) with flags “-c -s 20”^67,69^. DSD evaluates inter-gene network distance based on differences of diffusion by random walks from the two genes, and was a major portion of the best overall method for network module detection in Choobdar *et al*^100^. A subset of the DSD distance matrix containing only baits was used as input to the R function hclust with agglomeration method set to “ward.D2”. The resultant hierarchical clustering was divided into modules by the function cutreeHybrid in R package dynamicTreeCut (version 1.63-1) with parameters minClusterSize = 7, deepSplit = 2, method = ’hybrid’.

