## supplemental figure legend for "Autism genes converge on microtubule biology and RNA-binding proteins during excitatory neurogenesis"

**Supplemental figure legends:**

**Figure S1 Characterization of gene expression and cell cycle phase occupancy in NPCs and neurons**

(A) We quantified the protein expression of SOX2, a neural progenitor cell marker, and Ki67, a proliferation marker in NPCs. Percentage of SOX2 positive (SOX2+) cells and Ki67 positive (Ki67+) cells were measured by flow cytometry.

(B) During a 22-day neural differentiation, transcript expression for neural progenitors, differentiation, layer-specific, and proliferation markers were measured by bulk transcriptome profiling.Duplicate samples were collected at day 0, day 7, day 14 and day 22 of differentiation. The mean of normalized counts per million (cpm) for each gene from the duplicates was plotted.

(C) In the same 22-day differentiation experiment, the mean cpm across duplicates for each of the hcASD gene was plotted at each timepoint. 99 out of the 102 hcASD genes had non-zero count measured by bulk transcriptome profiling.

(D) Pie charts show the quartile analysis of the 99 hcASD genes at day 0, day 7, day 14 and day 22 of differentiation.

(E) Cell cycle phase occupancy measured by flow cytometry at d0, d2, and d4 differentiation of NPCs carrying a non-targeting control sgRNA. DNA content was labeled with 7-AAD and S phase cells were labeled with EdU.

**Figure S2 On-target knockdown efficiency of hcASD sgRNAs, pseudotime analysis and cluster mapping of cells from the CROPseq experiment**

(A) Log2FC of 73 hcASD genes show the on-target knockdown efficiencies. ▼denotes insignificant knockdown of the target gene and **x** denotes the target gene expression is below the detection threshold in our CROP-Seq experiment.

(B) Nine Leiden clusters were generated from all cells across 3 timepoints and RNA-velocity analysis predicted differentiation trajectories.

(C) Pseudotime analysis of all cells across 3 timepoints.

(D) A comparison between the target gene knockdown levels (log2FC) of 15 out of the 17 $ASD_{ps}$ genes and 66 out of the remaining 70 hcASD genes. Two sided t-test was performed.

(E) A comparison between the FDRs (Satterstrom *et al.*, 2020) of the 17 $ASD_{ps}$ genes and the remaining of the 70 hcASD genes. Two sided t-test was performed.

(F) The cumulative density of pseudotime for individual knockdowns showed unique trajectories of pseudotime and how they deviated at specific points from the non-targeting controls (NTCs). Decreased cumulative density from NTCs at earlier pseudotime indicates acceleration in pseudotime (*SUV420H1*, *KIAA0232*) whereas increased cumulative density from NTCs at earlier pseudotime indicates deceleration in pseudotime (*AP2S1*, *NSD1*).

(G) Cluster mapping using scRNA-seq data from prenatal cortical tissues[1] indicates that cells from our CROP-Seq experiment are mapped to the developing excitatory lineage in the human cortex. Top panel: 9 leiden clusters identified in our CROP-Seq experiment.

Middle panel: mapping cells using cell type marker genes from an external dataset. Bottom panel: mapping clusters using cell type marker genes from an external dataset.

(H) Cluster mapping using scRNA-seq data from developing human telencephalon[2] suggests that cells from our CROP-Seq experiment exhibit features similar to various stages of cells during differentiation of dividing intermediate progenitor cells (IPCs) toward excitatory neurons in the prefrontal cortex. Top panel: 9 leiden clusters identified in our CROP-Seq experiment. Middle panel: mapping cells using cell type marker genes from an external dataset. Bottom panel: mapping clusters using cell type marker genes from an external dataset.

**Figure S3 Experimental validations of cellular phenotypes in NPCs and neurons with $ASD_{ps}$ gene knockdowns**

(A) Knockdown efficiencies of the 17 $ASD_{ps}$ gene sgRNAs were evaluated in individually established NPC lines by qPCR. One sided t-test was performed. *$p < 0.05$, **$p < 0.01$, ***$p < 0.001$, ****$p < 0.0001$.

(B) Heatmap displays log2FC of cell cycle phase occupancies in G1, S, and G2/M of $ASD_{ps}$ gene knocked down cells over control cells for NPCs and day 2 neurons. One-way ANOVA was performed. *adjusted $p < 0.05$, **adjusted $p < 0.01$, *** adjusted $p < 0.001$, **** adjusted $p < 0.0001$.

(C) Representative images of immunocytochemistry show expression of a pan-neuronal marker TUBB3 and a deep layer neuron marker TBR1 in day 4 neurons containing non-targeting control sgRNA (NT-sgRNA) and hcASD sgRNAs targeting $ASD_{dc}$ gene *AP2S1* and *NSD1*, and $ASD_{ac}$ genes *KIAA0232* and *SUV420H1*. Experiments were performed in two batches and fluorescence intensity of TUBB3 and TBR1 of hcASD gene knocked down cells were compared to control cells in their own batch. Scale bar 30um.

(D) Percentage of cleaved caspase 3 positive (CCP3+) cells were quantified by flow cytometry for non-targeting control cells and *AP2S1*, *NSD1*, *KIAA0232* and *SUV420H1* knocked down cells at day 0, day 2, and day 4 of differentiation. Percentages of CCP3+ cells of hcASD gene knockdowns were normalized to matched non-targeting control cells as log2FC.

**Figure S4 sgRNA representation analysis and gene set enrichment analysis (GSEA) of the *Cycling 2* cluster.**

(A-C) Cluster shift of sgRNA representation at day 0 (A), day 2 (B), and day 4 (C) of differentiation. hcASD gene knockdowns that had at least one significant cluster shift at either of the timepoints: global (cells from all three timepoints combined together; Figure 5A), day 0, day 2 or day 4 were included in each heatmap.

(D) Heatmap shows change of sgRNA counts by timepoint. Counts of non-targeting controls (NTCs) and hcASD sgRNAs at day 2 and day 4 of differentiation were normalized to their counts at day 0 as fold change. The z-score of fold change was plotted for each of the 87 hcASD sgRNAs compared to NTCs.

(E) IDEA plot shows the significantly enriched molecular signatures (MSigDB Hallmark, 2023) of the top 200 upregulated genes of *Cycling 2* cluster versus the other three macro-clusters.
(F) IDEA plot shows the significantly enriched GO Biological Process terms of the top 200 upregulated genes of *Cycling 2* cluster versus *Cycling 1* cluster.
(G) IDEA plot shows the significantly enriched GO Biological Process terms of the top 200 upregulated genes of the *Cycling 2* cluster versus *Intermediate* cluster.
(H) Enrichment of *HNRNPH1* in *Cycling 2* cluster. Top panel: the 4 macro-clusters we identified in the CROP-Seq experiment (Figure 3E). Bottom panel: visualization of *HNRNPH1* expression on the UMAP.

**Figure S5 Differentially expressed gene analysis across hcASD gene knockdowns**
(A) Bar plot shows the effect size for 73 hcASD gene knockdowns as the number of differentially expressed genes (DEGs). Knockdown of the target gene itself is included in the counts.
(B) Distribution of the 3175 DEGs across all 87 hcASD gene knockdowns. Total occurrence represents the number of knockdowns a DEG belongs to.
(C) A comparison between the number of DEGs of the 17 $ASD_{ps}$ genes and the rest of the 70 hcASD genes. Two sided wilcoxon rank sum test was performed. ****p value < 0.0001.
(D) Distribution of the 2527 DEGs across the 17 $ASD_{ps}$ gene knockdowns. Total occurrence represents the number of knockdowns a DEG belongs to.
(E) Pairwise comparison of log2FC of the 2527 DEGs across the 17 $ASD_{ps}$ genes. Colored boxes of the dendrogram on the left indicate $ASD_{ac}$ (red) and $ASD_{dc}$ genes (green).
(F) IDEA plot shows significantly enriched molecular signatures of the 286 DEGs using MSigDB Hallmark (2023).

**Figure S6: Convergent differential expression profiles of RNA binding proteins.**
(A) Overview of methodology to identify clusters within ASD-PPI that are more densely interconnected.
(B) ASD-PPI divides into 8 clusters.
(C) CRISPRi knockdown of hcASD genes reveals targeted effects on individual hnRNP family members. hnRNP genes were grouped by family on the X-axis. The color of the cell corresponds to the “signed” -log10 of the FDR for differential expression downstream of perturbation of a specific hcASD gene (Y-axis), with purple indicating downregulation and green indicating upregulation.
(D) Functional network analysis of hcASD knockdowns highlights regulated *HNRNP* targets and their direction of change.
