## Supplementary figures and images for "Autism genes converge on microtubule biology and RNA-binding proteins during excitatory neurogenesis"

### Figure_S1

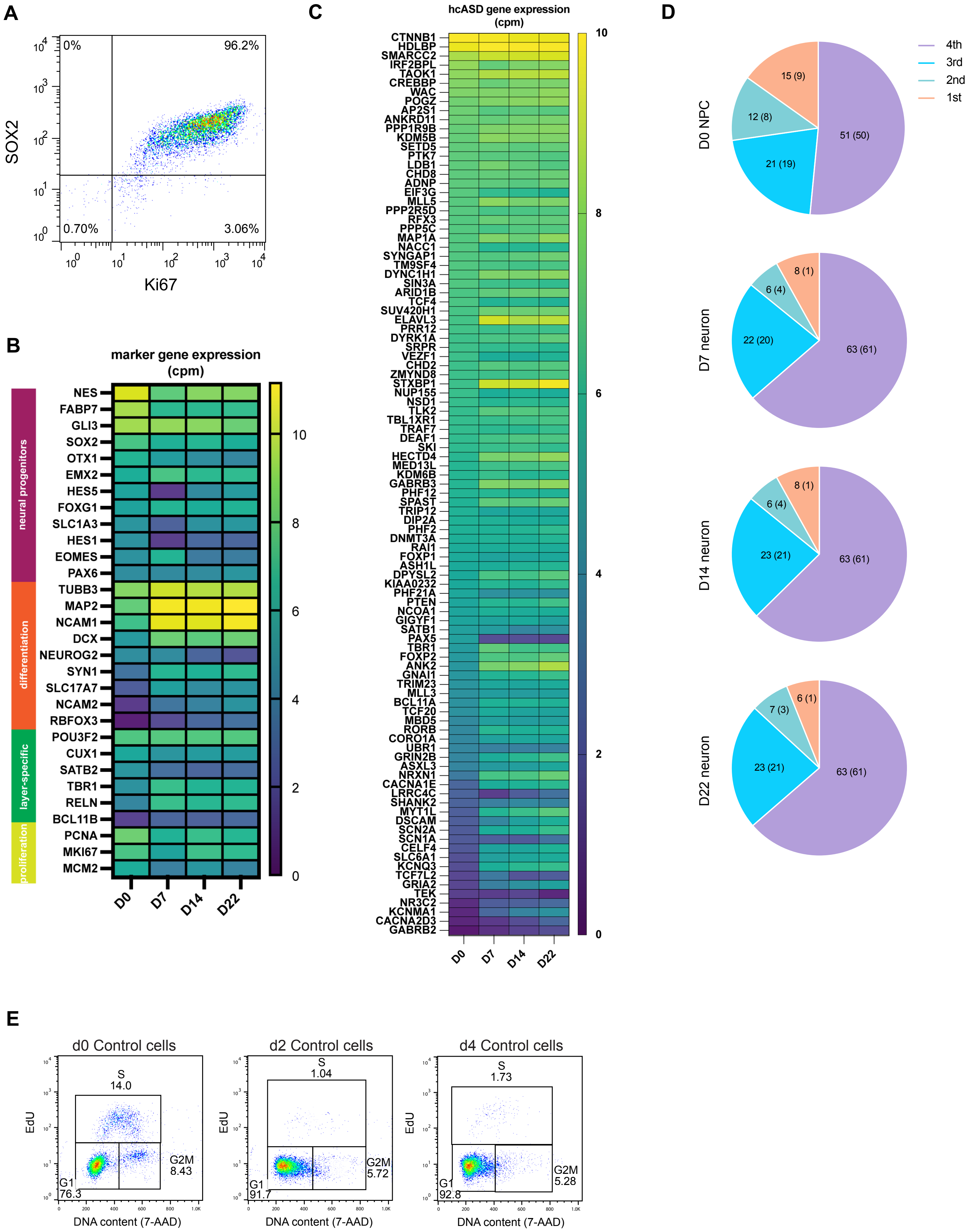

### Figure_S2

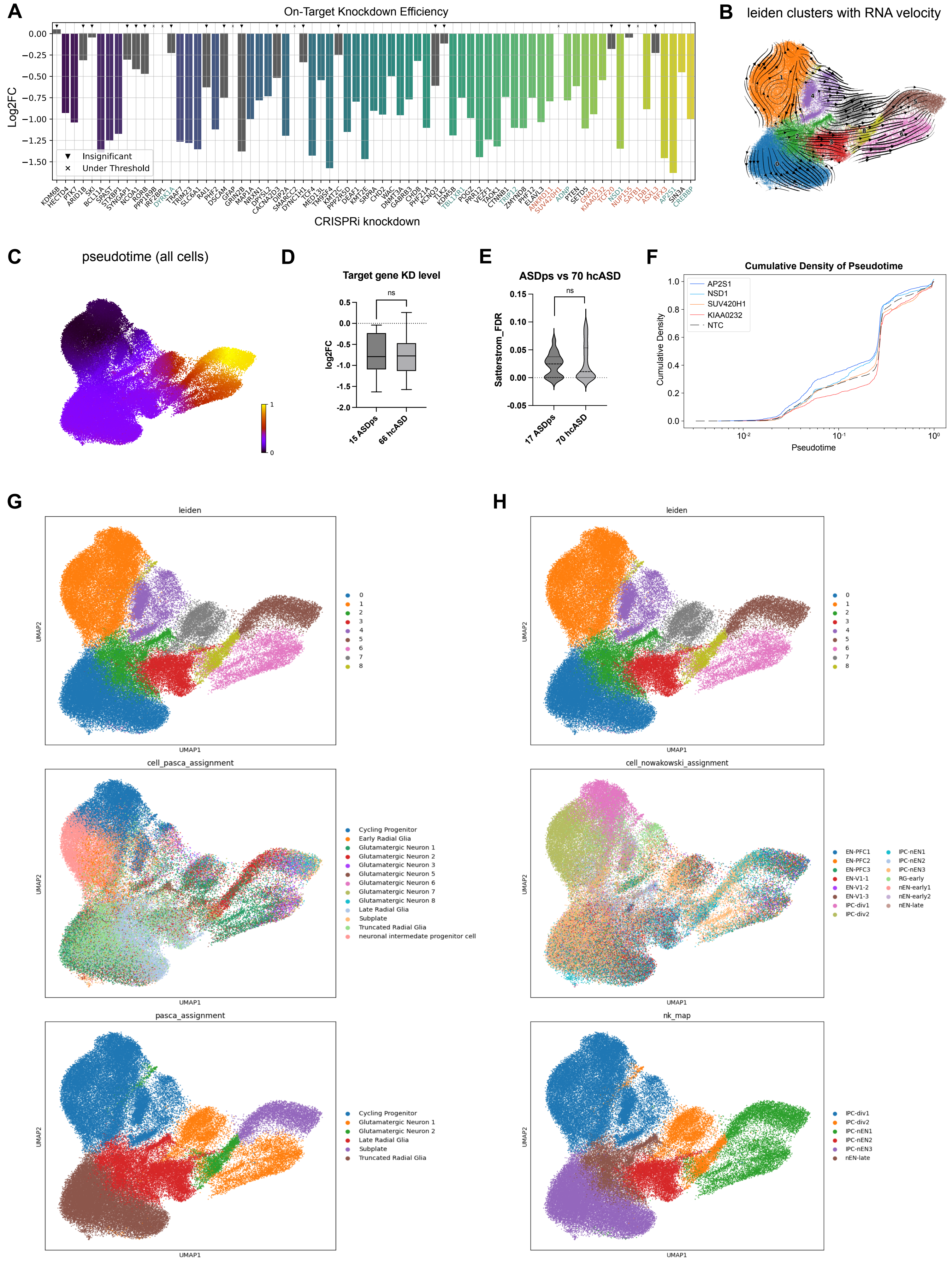

### Figure_S3

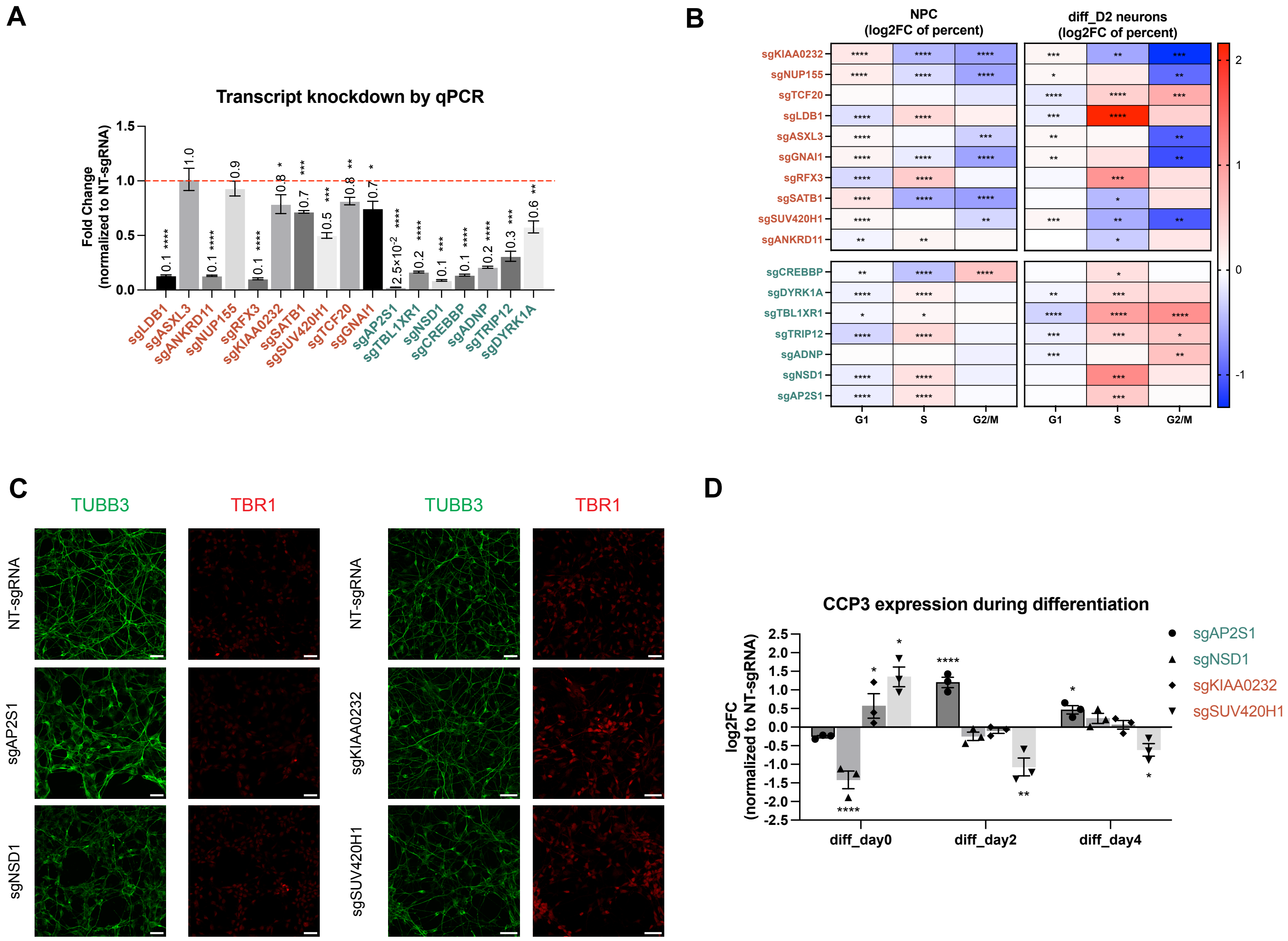

### Figure_S4

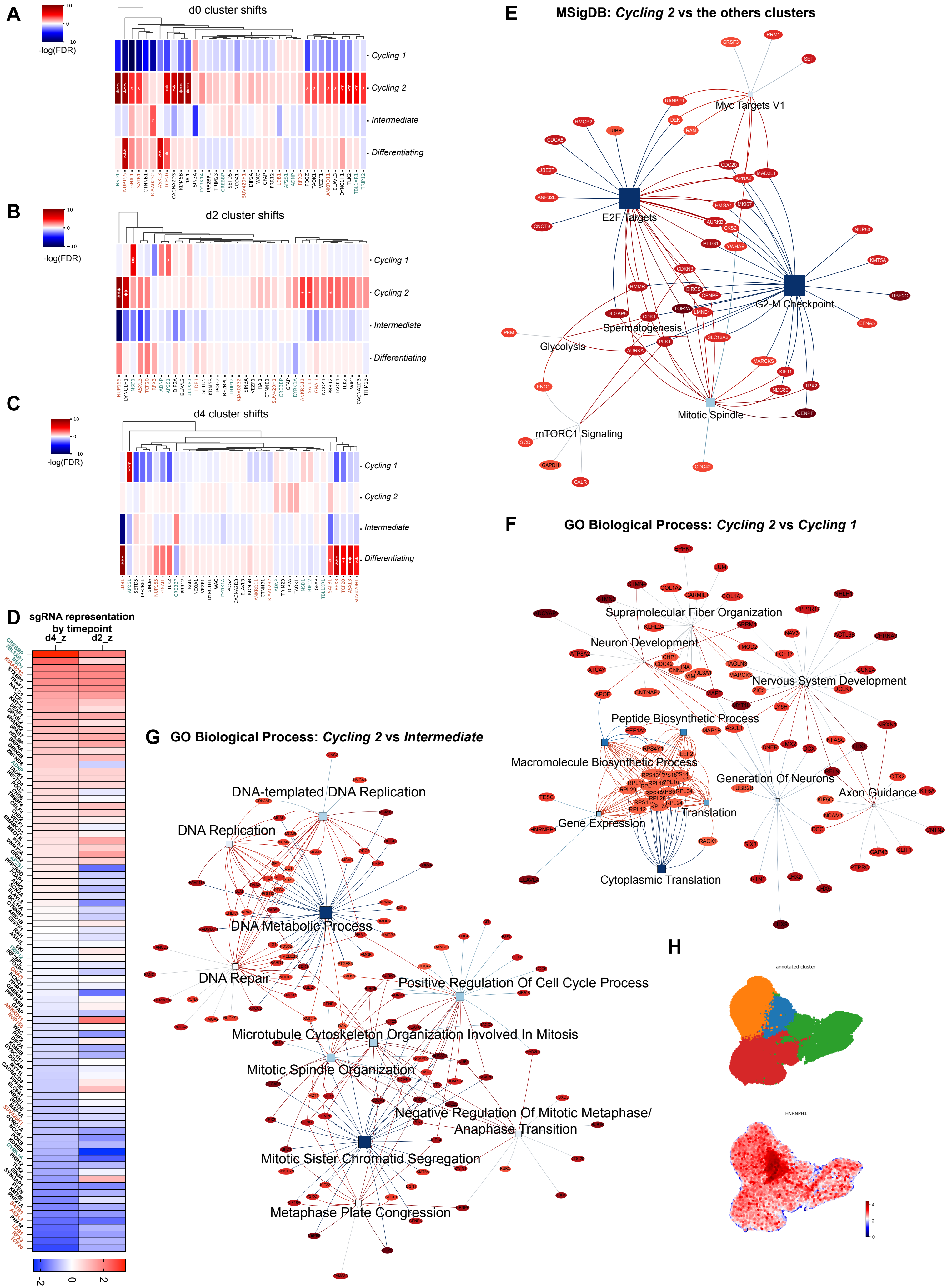

### Figure_S5

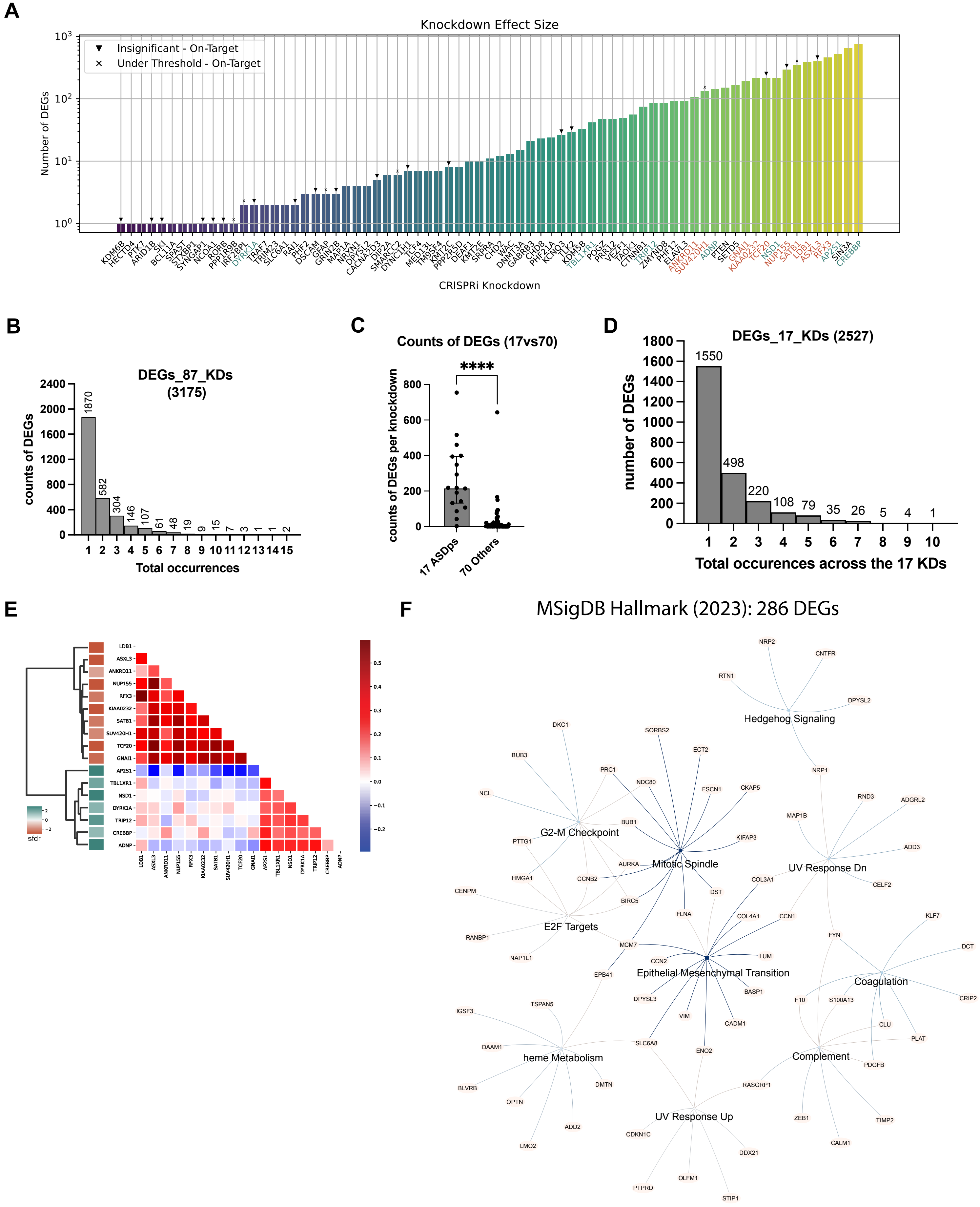

### Figure_S6

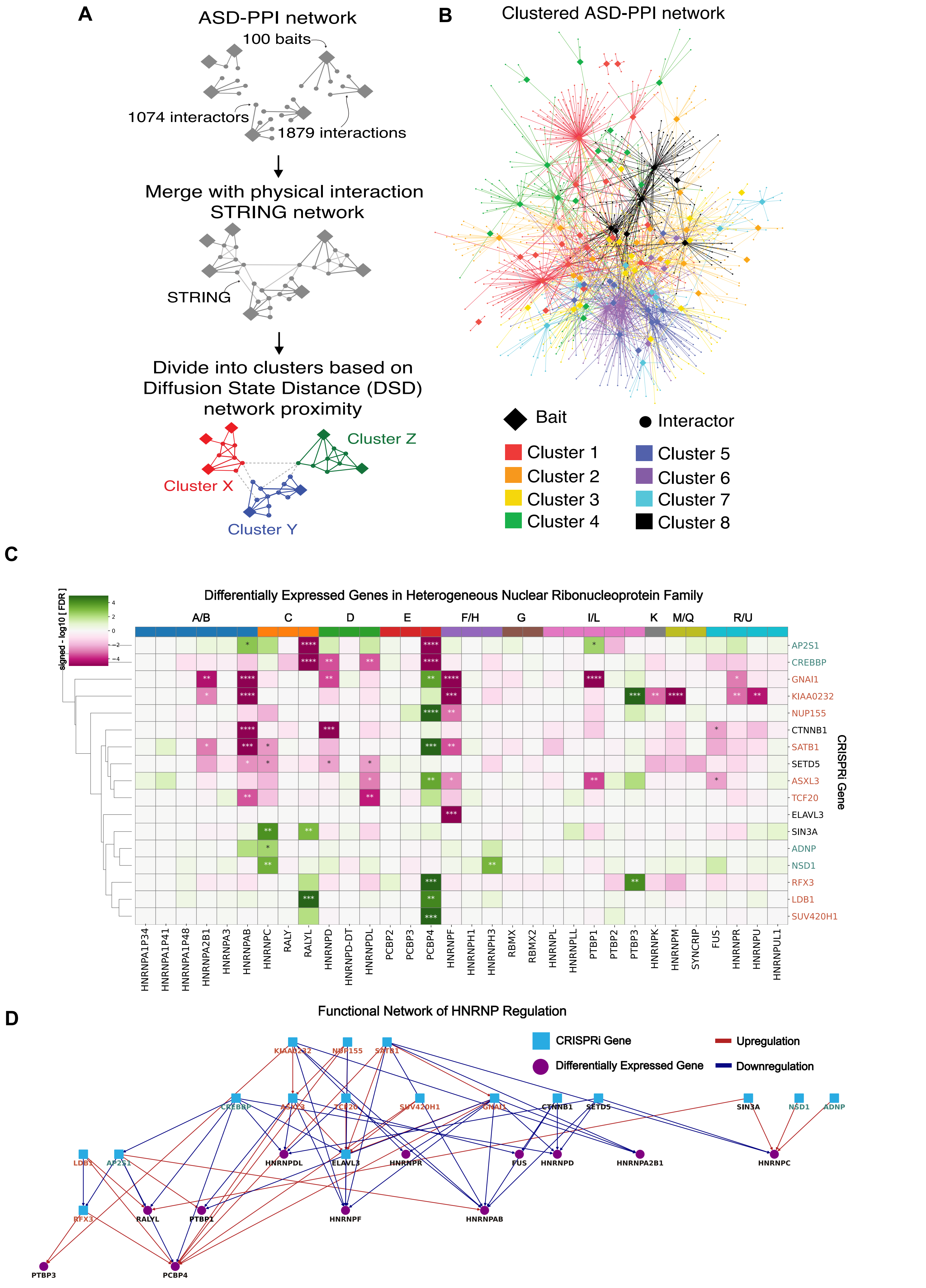
